# Elevated body temperature exacerbates arrhythmia and seizure-like activity in a zebrafish model of Timothy syndrome

**DOI:** 10.1101/2025.03.11.642683

**Authors:** Svetlana A. Semenova, Amir M. Sams, Deepthi Nammi, Matthew Menold, Gennady Margolin, Jennifer L. Sinclair, Katelyn A. Robertson, Kerry Larkin, Sydney Kelly, Andy Golden, Benjamin Feldman, Harold A. Burgess

## Abstract

Timothy syndrome (TS) is a multisystem disorder with autistic-like features, seizures and arrhythmia as the main symptoms. Most TS cases are caused by a *de novo* single amino acid substitution G406R in the *CACNA1C* gene that encodes the pore-forming subunit of the voltage-gated L-type calcium channel Ca_V_1.2. We generated a zebrafish model of TS with a homologous amino acid substitution in the *cacna1c*-encoded protein. Unlike patients, heterozygous mutants showed only mild impairments with no changes in mortality. However, homozygous mutants showed increased mortality, arrhythmias, neural activity and sensitivity to pentylenetetrazole-induced seizure-like behavior. Mutants also exhibited microcephaly, cerebellar hypotrophy and abnormal development of GABAergic neuron populations, and transcriptomic analysis revealed dysregulated expression of neuropeptide genes (including *bdnf* and *vgf*). Consistent with the idea that Ca_V_1.2 channels activate during fever, we found that heterozygous larvae manifested arrhythmia and seizure-like behavior when exposed to elevated environmental temperature, and homozygous larvae switched from bradycardia to tachycardia. These data provide a basis for using zebrafish to study the etiology of TS abnormalities and suggest t hat fever may be a particular risk for TS patients.

## Introduction

Timothy syndrome (TS) is a rare autosomal dominant disorder caused by *de novo* gain of function mutations in *CACNA1C*, the gene encoding the pore-forming α-1 subunit of the L-type voltage-gated calcium channel (LTCC) Ca_V_1.2. Major symptoms of TS include cardiac arrhythmia with QT interval prolongation, cutaneous syndactyly and autistic behaviors (reviewed in (*1*)). Most patients have a G406R missense mutation in either of two mutually exclusive, alternatively spliced isoforms of exon 8; mutations in exon 8A cause Timothy Syndrome 1 (TS1), whereas mutations in exon 8 cause Timothy Syndrome 2 (TS2). Each exon 8 variant encodes 34 amino acids with partially similar amino acid sequences, and channels with mutations in either exon 8 or 8A show sustained depolarization due to a reduction in voltage-dependent inactivation (*2*).

*CACNA1C* is expressed widely at multiple stages and in several cell types during neural development. Early expression of the exon 8A isoform occurs in neural progenitors, with a switch to the exon 8 isoform as neurons begin to differentiate (*2, 3*). Autistic symptoms in TS patients are likely due to changes in neural development, but the precise link is unclear, and results from studies to date have not been entirely consistent. In a hypomorphic mouse TS2 knock-in model, heterozygous animals showed impaired social interactions and an increase in repetitive behavior (*4*). These phenotypes were accompanied by an increase in migration of inhibitory neurons into the developing cortex, and an excess of inhibitory synapses (*5*). In contrast, in human brain organoids formed by patient-derived induced pluripotent stem cells (iPSCs), inhibitory neuron migration into the cortex was disrupted (*6–8*). Excitatory cortical neurons were also affected by increased *CACNA1C* channel activity, with concordant fate changes during differentiation of patient iPSCs and after overexpression of *CACNA1C* in embryonic mouse brains *in utero* (*2*). Although most studies have focused on cortical defects, a TS2 mouse model demonstrated abnormal activity in *CACNA1C* expressing hypothalamic glucose-sensing neurons that may relate to patient hypoglycemia (*9*). Thus, despite the widespread expression of *CACNA1C* during brain development, studies to date have characterized the effect of mutations on only a few brain regions.

In zebrafish, neural development can be visualized in real-time in the intact embryo and effects of mutations studied systematically using cellular resolution whole-brain imaging (*10, 11*). A zebrafish *cacna1c* loss of function allele caused developmental defects in ventricular cardiomyocytes and lower jaw development; however gain of function phenotypes relevant to Timothy syndrome, and effects on brain structure and function have not been examined (*12, 13*). We therefore generated a zebrafish model with a mutation homologous to the TS2 exon 8 G406R mutation, compared phenotypes to clinical manifestations, and used whole-brain imaging to assess changes in neural architecture.

Homozygous TS2 mutants showed a range of phenotypes reminiscent of clinical manifestations in patients. At embryonic stages, both cardiac chambers in TS2 homozygotes showed reduced contraction frequency, reminiscent of bradycardia reported at prenatal and early postnatal stages in patients (*14*). Several days later, contraction frequency normalized but mutants demonstrated an intermittent 2:1 atrioventricular block, recapitulating a characteristic patient arrhythmia. Epilepsy is present in a high proportion of patients, and accordingly, TS2 homozygotes showed heightened sensitivity to pentylenetetrazol, a seizure-inducing agent. Transcriptomic analysis revealed significant changes in the expression of genes encoding neuropeptides and peptide hormones in mutant larvae, with intermediate changes in heterozygotes. We used brain-wide morphometry to systematically assess changes in brain structure and composition. Mutants showed mild microcephaly and a disproportionately severe reduction in cerebellar tissue adjacent to the midbrain-hindbrain boundary (MHB). Differentiated neuron markers were inappropriately present at the midline in mutants, and inhibitory neuron markers were altered in a region-specific manner. Unlike patients, who are affected by dominant mutations in *CACNA1C*, heterozygous larvae did not manifest cardiac abnormalities or seizure-like behavior but did show subtle MHB hypotrophy, abnormal mature neuron positioning and intermediate gene expression changes. However, when tested at elevated temperature, seizures and arrhythmias were evident in heterozygous larvae and were exacerbated in homozygotes. This is consistent with the established role for fever in aggravating other long QT syndromes, and the sensitivity of LTCCs to elevated body temperature (*15, 16*). These findings establish a new model for TS2, reveal changes in the structure and composition of the midbrain-hindbrain region in TS2 zebrafish and raise concerns about fever as a safety risk in TS patients.

## Results

### Cerebellar Purkinje cells gradually become the highest in cacna1c transcripts in the developing brain

Previous studies revealed that *cacna1c* is expressed during zebrafish embryonic development in heart, pancreas and jaw cartilage (*12, 13*). Single-cell sequencing data from Daniocell, a publicly available dataset generated from staged zebrafish embryos, showed that *cacna1c* expression increases in developing brain starting at around 36 hpf, closely paralleling the expression of *synaptophysin a* (*sypa*), a marker of mature neurons (Supp. Fig. 1)(*17*). We used fluorescent *in situ* hybridization to examine the distribution of *cacna1c* in developing brain. At 2 dpf, *cacna1c* expression was detected in the telencephalon, optic tectum, thalamus and rhombencephalon, although not adjacent to midline proliferative zones (Fig. 1A). *cacna1c*-expressing cells in the hindbrain formed bands parallel to the axis of the brain, which were especially distinct in coronal views (Fig. 1A’). At 3 dpf, expression in the telencephalon, optic tectum, thalamus and rhombencephalon persisted, and the area adjacent to the midline of the brain remained negative (Fig. 1B). Cell groups with especially high expression of *cacna1c* were found in the cerebellum, where an emerging cluster was visible. At 6 dpf, strong *cacna1c* continued to show widespread expression, including in the pallium, subpallium, diencephalon and optic tectum (Fig. 1C), with a stronger signal in pallium, subpallium and cerebellum (Fig. 1D-E). There were two strong distinct bilateral patches of expression within the cerebellum: a rostromedial cluster located in the teleost-specific valvula cerebelli and a caudolateral cluster that overlapped with co-registered scans for *gad1b*, a marker for inhibitory neurons (Fig. 1E2-3), and *pvalb6*, a gene that is highly expressed in mature Purkinje neurons (Fig. 1E4) (*18*). In contrast, cerebellar clusters enriched in *cacna1c* expression did not overlap with *neurod1*, a marker for granule cells (not shown). Thus, while *cacna1c* is expressed throughout the cerebellum there is marked enrichment in the valvula cerebelli and in Purkinje neurons. Dual *in situ* hybridization with *elavl3*, a gene that is most strongly expressed immediately as neurons become post-mitotic, then decreases to intermediate levels in mature neurons, revealed that in 3 dpf and 6 dpf brains, *cacna1c* did not overlap with the strongest areas of *elavl3* expression, but rather with the adjacent areas where *elavl3* was less robustly expressed (Fig. 1B inset and 1F-G). Thus, in the developing zebrafish brain, *cacna1c* is most strongly expressed in mature neurons rather than in proliferating cells or newly generated neurons.

**Figure 1.**
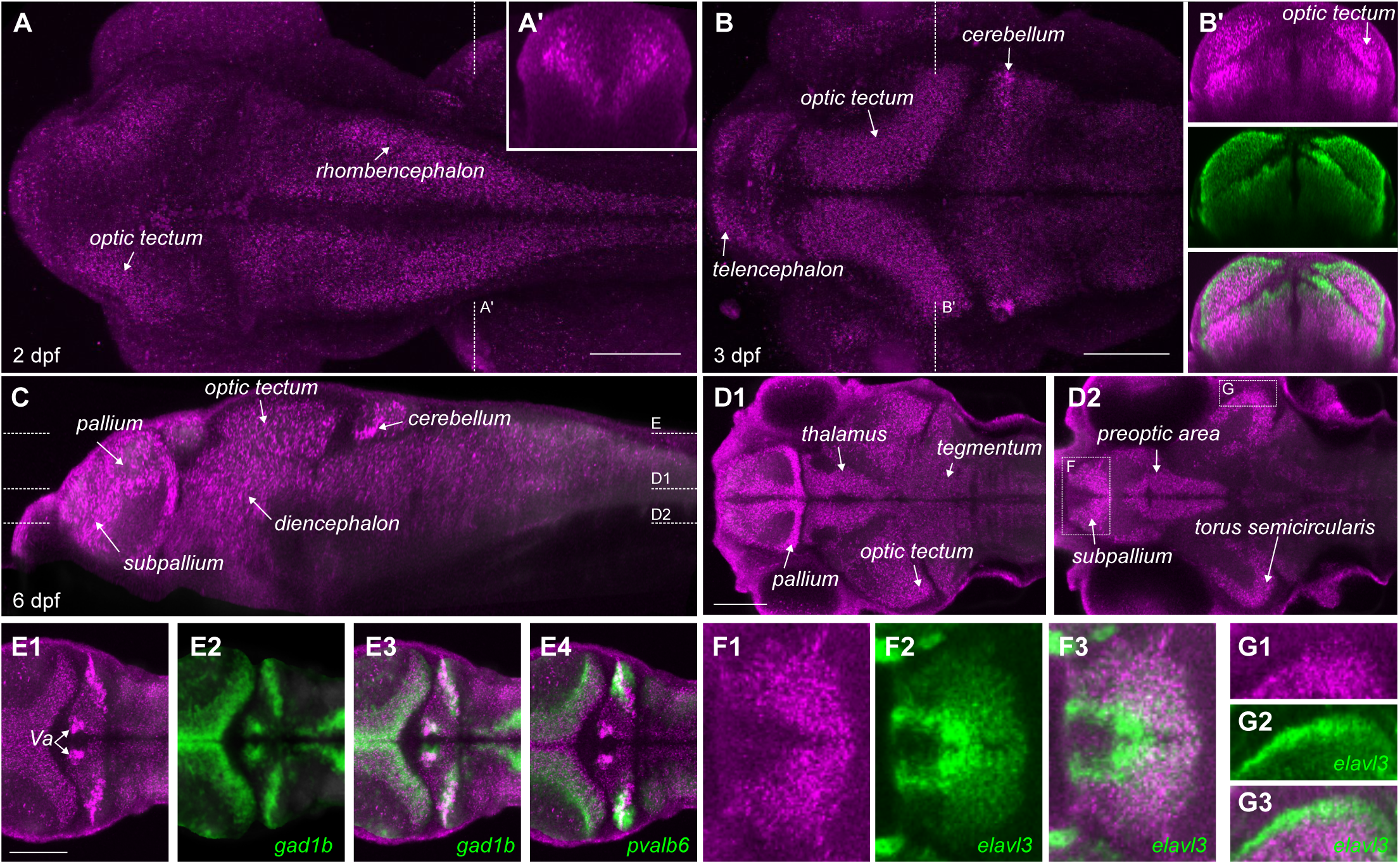
Enriched cacna1c expression in cerebellar neurons during neural development. A-B. Maximum projection of confocal stacks of *cacna1c* expression from *in situ* HCR at 2 dpf (A) and 3 dpf (B). Insets show coronal re-section at the level indicated by the dotted lines. In (B), inset shows co-*in situ* hybridization for *elavl3* (green). Scale bars 100 µm. C. Sagittal re-section of confocal image stack showing *cacna1c* expression at 6 dpf (mean of 5 co-registered brain images), with dotted lines indicating levels of horizontal slices in D and E. D. Horizontal confocal sections showing *cacna1c* expression in midbrain and forebrain. Scale bar 100 µm. E. Horizontal confocal section through the cerebellum, showing co-registered expression of *cacna1c* (magenta) with *gad1b* (E2-3, green) and *pvalb6* (E4, green)). Scale bar 100 µm. F-G. Horizontal section showing expression of *cacna1c* (magenta) relative to co-labeling with *elavl3* (green) in the subpallium (F) and torus semicircularis (G).

### Generation of zebrafish with Timothy syndrome 2 mutation

Human and zebrafish Cacna1c protein sequences are highly conserved, with 78.9% identity. We examined genomic sequence in zebrafish to assess the feasibility of generating a zebrafish mutation that precisely recapitulated patient mutations in Timothy syndrome. The G406R mutation in Timothy syndrome is due to a missense mutation that can occur at the end of either of the two mutually exclusive exon 8 variants, exon 8 or exon 8A. Patient mutations that affect either exon 8 or 8A have overlapping clinical manifestations, including autism, seizures and cardiac arrhythmia. However, mutations in exon 8 lead to worse outcomes than 8A mutations, and present other differences such as a lack of syndactyly (*19*). These phenotypic variations may be because exon 8 is around 4-fold more abundant in brain and heart tissue, or because exons 8 and 8A show distinct temporal patterns of expression during neural development. Similarly, the zebrafish homolog of *cacna1c* comprises alternate exons 8 and 8A, which encode amino acid sequences that are 100% and 94% identical to human exons 8 and 8A respectively (Fig. 2A). We performed qPCR at 2, 4 and 6 dpf to measure the relative expression of isoforms incorporating exons 8 and 8A in zebrafish. We did not detect an age-related change in exon usage (F_2,10_ = 2.89, p = 0.11), and exon 8 was at least 13-fold more abundant than exon 8A at all ages (Fig. 2B).

**Figure 2.**
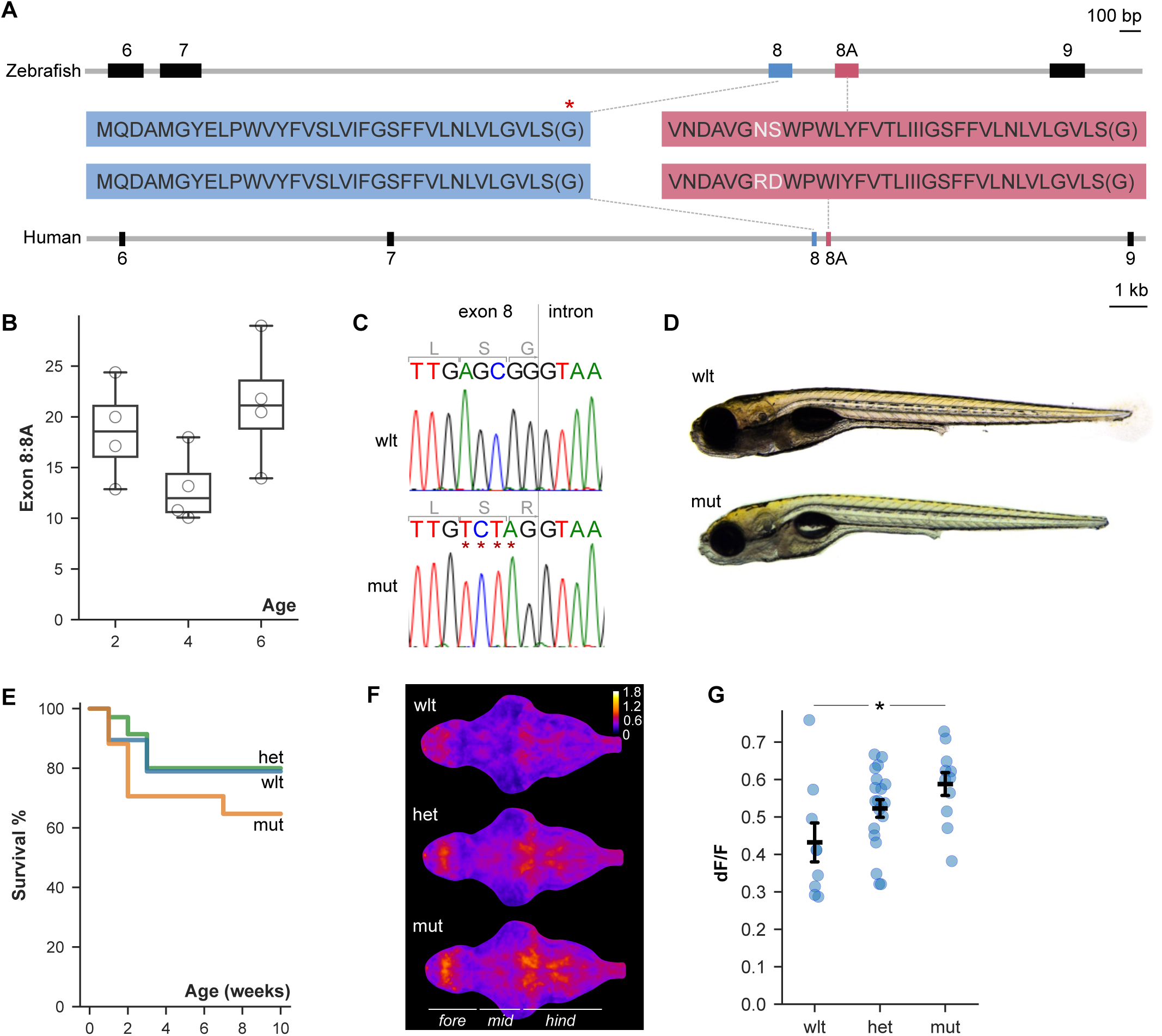
A zebrafish model for Timothy syndrome 2. A. Mutually exclusive exons 8 and 8A are conserved in zebrafish cacna1c and human CACNA1C genes. The codon for the glycine residue mutated in TS is shown in parentheses because its third base is contributed by exon 9. Red asterisk, location of the mutation in TS2. B. Ratio of exon 8 to 8A expression in 2, 4 and 6 dpf larvae (qPCR). C. Sequence from wild-type (top) and mutant (bottom) larvae. Red asterisk indicates the nucleotides edited (mutation that converts the final codon in exon 8 from encoding Gly to Arg and a silent mutation in the penultimate codon for Ser in exon 8, AGC to TCT, introduced to facilitate genotyping). D. TS2 wild-type and homozygous mutant larvae at 6 dpf. E. Kaplan Meier survival plot for wild-type, heterozygous and mutant larvae during development to adulthood (N=19,35,17 larvae respectively). F. Mean GCaMP dF/F in 6 dpf wild-type, heterozygous and mutant larval brains (N = 9, 20, 11 respectively). Forebrain, midbrain, hindbrain indicated. Color bar shows dF/F G. GCaMP dF/F within cellular brain area for samples in (G). Mean and standard error of mean indicated for each genotype. * p = 0.014.

We generated a G419R missense mutation (corresponding to human G406R) at the end of the more highly expressed exon 8, thereby creating allele *cacna1c^y680^* as a potential model for Timothy syndrome 2 (herein referred to as the TS2 mutant; Fig. 2C). Homozygous mutant (herein ‘mutant’) and heterozygous TS2 larvae were overtly similar to their wild-type siblings in body size and overall appearance (Fig. 2D). Early mortality is common in TS patients. To characterize the effect of *cacna1c* G419R homozygosity on survival, we genotyped embryos from a TS2 heterozygous incross then raised larvae in mixed genotype pools. Under these conditions, only 23% of mutants survived to adulthood, significantly less than the survival rates of 69% and 90% for heterozygous and wild-type fish respectively (ANOVA main effect of genotype, F_2,6_ = 25.037, p = 0.001; Supp. Fig. 2A). However, when raised in same-genotype pools, the mortality of mutants was only 18% greater than the mortality of wild-type siblings; diminished survival in mutants may therefore reflect a reduced ability to compete with siblings (Fig. 2E). Surviving mutants were sexually dimorphic and fertile, although males were smaller than wild-type siblings (ANOVA, interaction of genotype and sex, F_2,50_ = 3.5, p = 0.037; Supp. Fig. 2B). Because the G419R mutation resulted from nucleotide changes at the 3’ end of exon 8, we questioned whether the relatively normal growth and survival observed in mutants might be due to a splicing change that minimized the mutation’s effects by increasing expression of isoforms incorporating the wild-type exon 8A. However, we found no significant changes in the frequency of splicing into or out of either exon 8 or 8A in mutants, making it unlikely that splice changes mask the mutant phenotype (Supp. Fig. 2C-D). Thus, unlike in mammals where TS mutations are dominant and lead to high mortality, heterozygous TS2 zebrafish are overtly normal and even homozygotes are viable and fertile (*9*).

We anticipated that the TS2 mutation would lead to a sustained depolarization in *cacna1c-*expressing cells, causing elevated activity. To test this, we crossed the TS2 mutation to a pan-neuronally expressed GCaMP and measured fluorescence intensity over a 5 minute period for each larva. Mutant larvae showed greater changes in fluorescence compared to wild-types, confirming that the TS2 mutation leads to increased neuronal activity (F_2,37_ = 4.4, p = 0.019; Fig. 2F-G, Supp. Video 1).

### Increased expression of neuropeptides in TS2 mutants

Next we assessed changes in gene expression, using RNA-seq to compare wild-type, heterozygous and homozygous TS2 larvae at 2 and 6 dpf. We prepared total mRNA after removing the entire trunk caudal to the swim bladder to reduce the contribution of skeletal muscle genes and enrich for genes expressed in the brain and heart. At 2 dpf, 306 genes were differentially expressed in TS2 mutants compared to wild-types (149 and 157 down and upregulated respectively; Supp. Fig. 3B). Pathway analysis revealed significant upregulation of genes involved in muscle development (*actn3b, dysf, neb, ttn.1, ttn.2, lama2, obscnb, ctsd, smad4a, smyhc1*) and cardiac muscle contraction (*trdn, cacna1sb, actc1c, atp2a1l, ACTC1, atp1a1a.4*), and downregulation of genes involved in mRNA splicing (*srsf4, ddx5, snrpd3l, lsm4, bud31, snrpf, rbm8a, snrpd2, hsp70l*). At 6 dpf, 46 genes were significantly upregulated and 25 genes downregulated in mutants (Fig. 3A). Downregulated genes were involved in cell cycle (*bub1, mcm7, ttk, smad3a*), whereas upregulated genes were implicated in neuropeptide hormone signaling (*bdnf, vgf, si:dkey-175g6.2, npb, npy, adcyap1b*; Fig. 3B). Two other upregulated genes – *star* and *si:dkey-58b18.8* – are expressed in the anterior hypothalamus of larval zebrafish, where many peptidergic neurons are located (*20*).

**Figure 3.**
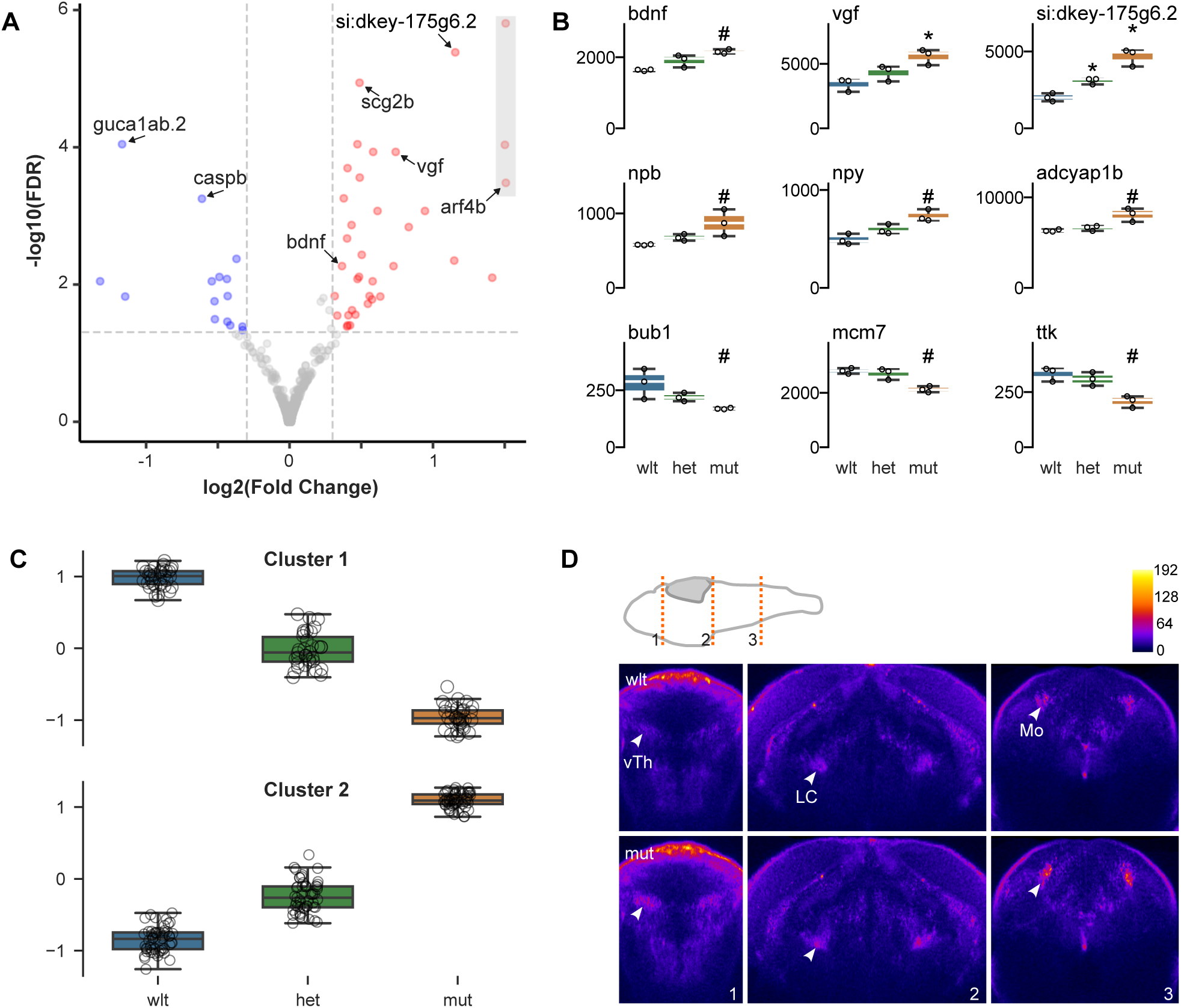
Changes in neuropeptide gene expression. A. Volcano plot for genes differentially expressed between wild-type and mutant 6 dpf larvae using RNA-seq from anterior tissue (includes head and heart). Values for points in grey rectangle truncated for display. B. Normalized read counts by genotype for selected differentially expressed genes. DESeq2 adjusted p-values: * p < 0.001. # p < 0.1 C. Normalized expression of genes present in the two largest clusters from the *ConcensusClustersPlus* clustering algorithm, each comprising genes with correlated levels of expression in wild-type, heterozygous and mutant larvae. Input genes were defined based on differential expression between wild-type and mutant larvae, revealing an intermediate gene expression phenotype in heterozygotes. D. Pseudocolored (see color scale bar) coronal sections through the diencephalon, pons and medulla after HCR fluorescent *in situ* hybridization for *vgf* in wild-type (top panels) and mutant (bottom) 6 dpf larvae. Images are the mean of multiple larvae that were co-registered (N = 9,8 for wild-type and mutant respectively). Schematic shows the position of each coronal plane. vTh, ventral thalamus; LC, locus coeruleus; Mo, medulla oblongata.

We also compared heterozygous mutants to wild-type siblings at 6 dpf. Only 10 genes were differentially expressed. For four of these, the unadjusted p-value for the wild-type/mutant comparison was also less than 0.05, and expression changed in the same direction (*fhl5, gcgb, smyhc1, si:dkey-175g6.2*; Supp Fig. 3C). To examine whether subtle changes were present in heterozygous larvae, we used the subset of genes that were differentially expressed between 6 dpf wild-type and homozygous mutants as input for the ConcensusClusterPlus algorithm to perform unsupervised clustering of genes into groups based on correlated expression in wild-type, heterozygous and mutant larvae. Most (95%) genes were assigned to two large clusters corresponding to genes that were downregulated and upregulated in mutants, respectively. Interestingly, mean expression levels in heterozygous animals were midway between wild-type and mutant levels in each cluster (Fig. 3C). Thus, although heterozygous larvae are overtly normal, they nonetheless manifest changes in gene expression that are similar to, but less severe than in homozygous mutants.

Interestingly, two of the neuropeptides that were upregulated in mutants (*vgf* and *si:dkey-175g6.2*) are paralogs of mammalian VGF, and are known to be positively regulated by a third upregulated gene, *bdnf*. Because VGF has been implicated in neural development in mice, we examined *vgf* expression in mutants using fluorescent *in situ* hybridization (*21, 22*). *vgf* was expressed in discrete clusters in the pallium, subpallium, ventral thalamus, raphe, locus coeruleus and medulla oblongata, with mutants showing increased expression compared to wild-type siblings in each of these areas (Fig. 3D).

### Embryonic bradycardia and intermittent cardiac arrhythmia in cacna1c mutants

Life-threatening episodes of arrhythmia (including long QT-interval, bradycardia and 2:1 atrioventricular block) are a characteristic feature of Timothy syndrome. In a loss-of-function *cacna1c* allele (*island beat*), atrial cardiomyocytes from mutant embryos showed uncoordinated contractions, whereas ventricular cardiomyocytes failed to contract, even in response to direct electrical stimulation (*12*). We therefore used high-speed video recordings to assess contractions in each cardiac chamber separately. Contraction frequency was normal in 2 dpf heterozygous embryos (Supp. Fig. 4A) but was significantly reduced in both chambers in homozygotes (Fig. 4A-B, Supp. Fig. 4A).

**Figure 4.**
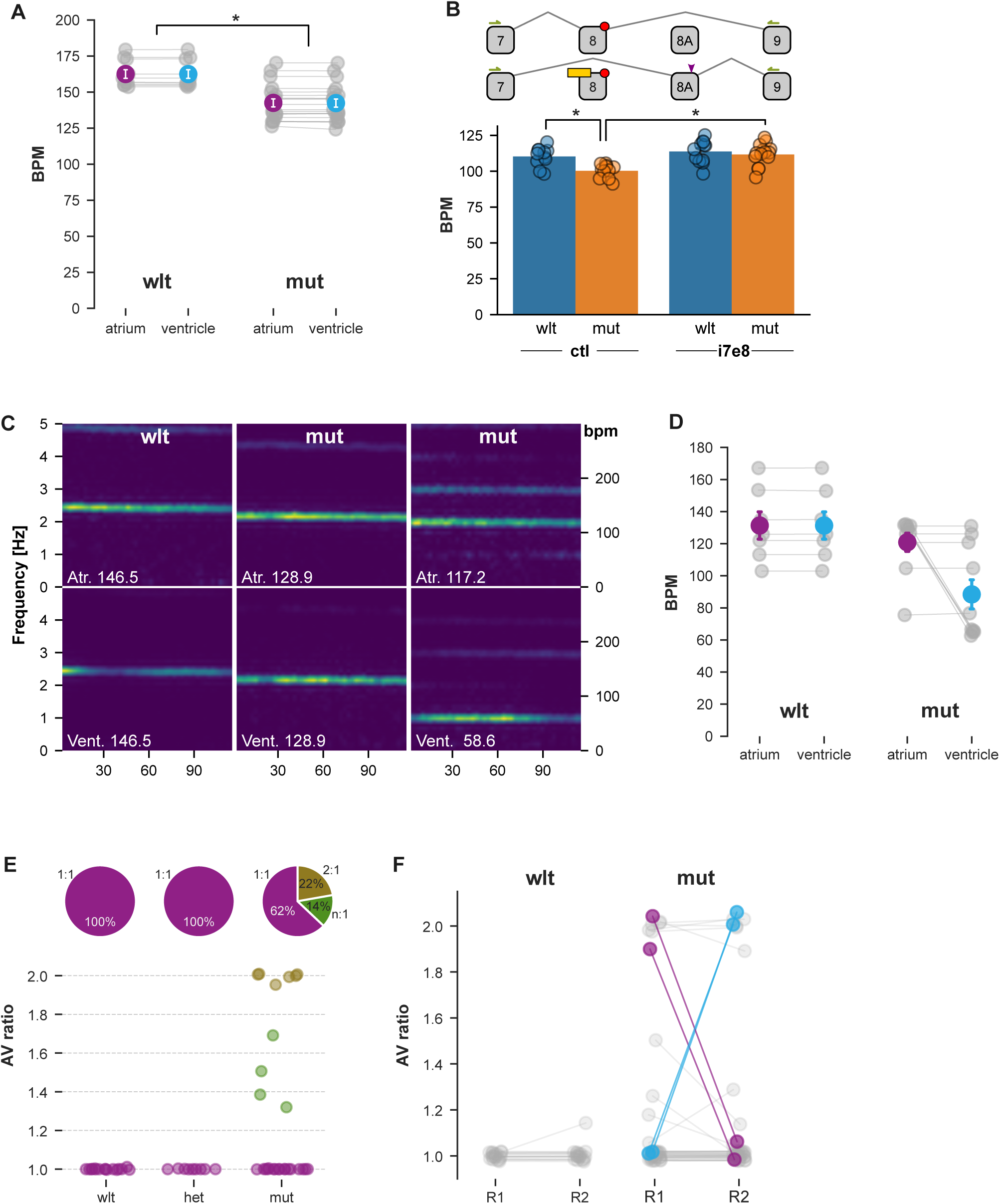
Embryonic bradycardia and intermittent arrhythmia phenotypes. A. Contraction frequency for cardiac chambers (beats per minute, bpm) at 2 dpf in wild-type and mutant TS2 larvae (N = 10, 20). * t-test p < 0.001 B. Antisense oligonucleotide (yellow) used to remove the exon 8 mutation (red) by shifting isoform expression from exon 8 to exon 8A. Arrows, RTPCR primers. Arrowhead, EcoRI site. Atrium contraction frequency at 2 dpf in wild-type and mutant larvae injected with standard control (ctl) or i7e8 splice-blocking morpholino (N = 7–10 / group). Mutants showed bradycardia (ANOVA, main effect of genotype, F_1,30_ = 6.6, p = 0.015). C. Spectrograms for 5 dpf wild-type (left panel) and mutants (middle, right panels) showing power at 0–5 Hz frequencies (right axis, 0–300 bpm). Top panels, atrium. Bottom panels, ventricle. D. Contraction frequencies in 5 dpf larvae (wild-type, N = 7; mutant, N = 10). Mean and standard error indicated. E. Atrium:Ventricle contraction frequency ratio (AV ratio) for wild-type, heterozygous and mutant larvae (N = 16, 9, 27). Piecharts: percent of larvae for each genotype with a 1:1, 2:1 or intermediate (n:1) ratio. F. AV ratio for wild-type and mutant (N = 17, 41) larvae recorded twice with at least a 4 hour interval between the first (R1) and second (R2) recording. Larvae that switched between AV ratios highlighted by colored lines.

The expression of alternative splice isoforms of *cacna1c* has been leveraged to correct deficits in a human TS1 organoid model (*8*). An antisense oligonucleotide that prevented splicing of exon 8A, which included the gain of function mutation, led to increased expression of the unaffected exon 8 and rescued deficits in neural development. We designed an antisense morpholino against the intron 7–exon 8 (i7e8) junction, which blocked splicing from exon 7 to exon 8, leading to increased inclusion of exon 8A (Supp. Fig. 4B). Wildtype and mutant embryos injected with the i7e8 morpholino showed increased cardiac contraction frequency, demonstrating a general effect of the morpholino (ANOVA main effect of treatment, F_1,44_ = 16.7, p < 0.001; Fig. 4B, Supp. Fig. 4C-D). As a result, mutants treated with the i7e8 morpholino showed a contraction frequency similar to wild-types injected with control morpholino and were therefore effectively normalized.

By 3 dpf, we noted that mean atrial contraction rates in homozygotes were no longer significantly reduced, but that in some larvae, the ventricle contraction frequency was half that of the atrium, a phenomenon seen in 2:1 atrioventricular (AV) block (Fig. 4C,D, Supp. Video 2). Whereas all wild-type and heterozygous larvae showed matched (1:1) AV contraction frequency, 22% (6/27) of mutants had a 2:1 rhythm, and 14% an intermediate ratio (Fig. 4E). To assess whether arrhythmia was a stable phenotype in individual mutant larvae, or whether arrhythmia was intermittent, we recorded from the same larvae twice, at an interval separated by 4-6 hours. Around 10% (4/41) of mutants switched between matched AV rhythms and a 2:1 beat frequency (Fig. 4F). We observed an abrupt switch from a matched AV rhythm to a 2:1 rhythm during one recording (Supp. Fig. 4E). Thus, TS2 homozygous mutants have bradycardia at early stages, later followed by intermittent cardiac arrhythmia, consistent with findings in Timothy syndrome patients.

### TS2 larvae display elevated seizure susceptibility

As seizure disorders are seen in a majority of TS2 patients (*14*), we assessed whether TS2 fish showed heightened seizure susceptibility. We exposed larvae to pentylenetetrazol (PTZ), a GABA receptor antagonist that, at low doses, only slightly elevates the risk of seizure-like activity in larval zebrafish (*23*). Larvae not exposed to PTZ did not exhibit genotype-related differences in locomotor activity (Fig. 5A-B, PTZ 0 mM). However, after PTZ treatment, TS2 mutants tended to show long periods of immobility that were broken by unusually vigorous swim bouts, similar to previous descriptions of seizure-like activity in zebrafish (Fig. 5B, PTZ 1 mM). The first reported TS2 patient was described as showing “frequent startle reflexes” (*19*), and a natural history study of a large patient cohort noted that loud noises often caused distress (*14*). While the exact cause of these auditory-associated symptoms in TS2 are unknown, we tested auditory sensory and sensorimotor gating but did not detect changes in escape reactivity or prepulse inhibition (Supp. Fig. 5A-C). Thus, TS2 zebrafish share seizure susceptibility with patients but not increased auditory reactivity.

**Figure 5.**
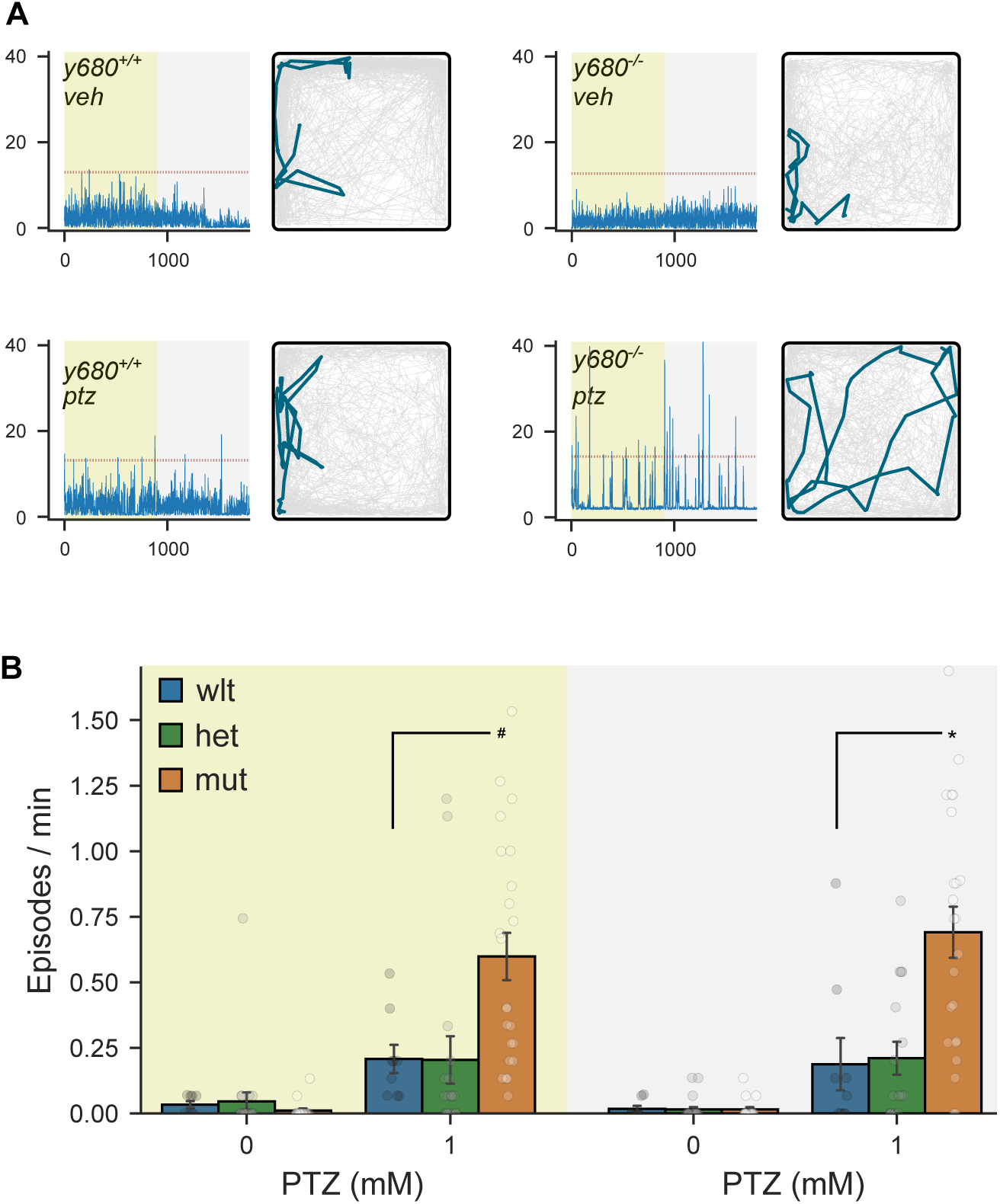
Increased seizure susceptibility after pentylenetetrazole treatment. A. Locomotor activity in individual TS2 wild-type and mutant larvae, under illuminated (yellow background) or dark (grey background) conditions, after exposure to 1 mM PTZ (ptz) or vehicle (veh). Red dotted line indicates distance threshold (total pixel movement in 1 sec interval) for designating seizure-like episodes. Traces to the right of each plot show the position of the larva over a 10 second interval during the light phase. B. Seizure-like movement episodes per minute in wild-type, heterozygote and mutant TS2 larvae recorded for 15 minutes in illuminated (left) or dark (right) conditions with vehicle alone or in 1 mM PTZ. # t-test p = 0.014 and * p = 0.006, for mutants compared to wild-types.

### Altered brain morphology in TS2 larvae

To systematically examine changes in brain structure in TS2 mutants, we measured brain structure and composition using voxel-based morphometry (*11*). For morphometric analysis, we imaged wild-type and mutant sibling larvae, each carrying two transgenes: the *tuba:mCardinal* transgene that labels all post-mitotic neurons, and the *gad1b:RFP* transgene that labels GABAergic neurons. We then used elastic registration to align confocal scans to the Zebrafish Brain Browser (ZBB) reference brain that was previously annotated with neuroanatomical labels and genetic markers (*24, 25*). Next, we computationally extracted clusters of significantly different voxels in the resulting transformation matrices to assess local changes in brain volume and clusters of voxels with altered fluorescence intensity to locate changes in *tuba:mCardinal* or *gad1b:RFP* signals (*11*).

Mean brain volume was similar in wild-type and heterozygous sibling 6 dpf larvae, but reduced by 6.1% in homozygous mutants (F_2,64_ = 4.3, p = 0.018; Supp. Fig. 6A). In homozygotes, there was a greater reduction in brain regions primarily occupied by neuronal somas than in primarily neuropil regions (F_2,65_ = 7.2, p = 0.001; Supp. Fig. 6B). We counted cell density in several brain regions in larvae stained with a nuclear dye, and found a significant reduction in the stratum periventriculare of the optic tectum, a cell dense region in the midbrain (Supp. Fig. 6C,D). Voxel-wise analysis revealed a salient cluster of voxels that were disproportionately compressed in a midline region in the anterior hindbrain, located between the cerebellar commissure and the caudal lobe of the optic tectum (Fig. 6A). This region is occupied neither by the somas of mature neurons nor by neuropil (Fig. 6B). To assess the reproducibility of this finding, we annotated the reference brain with a mask that covered the affected region, then imaged an independent cohort of larvae. The volume occupied by the mask was reduced by 8% in mutants compared to wild-types (even after normalizing for the smaller brain volume in mutants), confirming a change in brain architecture in this region (t-test, p < 0.001; Fig. 6C). Heterozygotes also showed 6.7% reduction in the same region (t-test, p < 0.001; Supp. Fig. 6E). The ZBB atlas revealed that the affected region is bounded by expression of *sox2*, a marker of neural stem cells, and overlaps with *en1b*, which labels the midbrain-hindbrain boundary (Fig. 6D,E). We performed *in situ* hybridization against *sox2* and *en1b* and found that the extent of *en1b* expression was reduced in mutants, overlapping with the region of reduced volume (Fig. 6F), whereas no changes in *sox2* expression were detected (Supp. Fig. 6F). Thus, the midbrain-hindbrain boundary region is disproportionately reduced in size in TS2 mutants.

**Figure 6.**
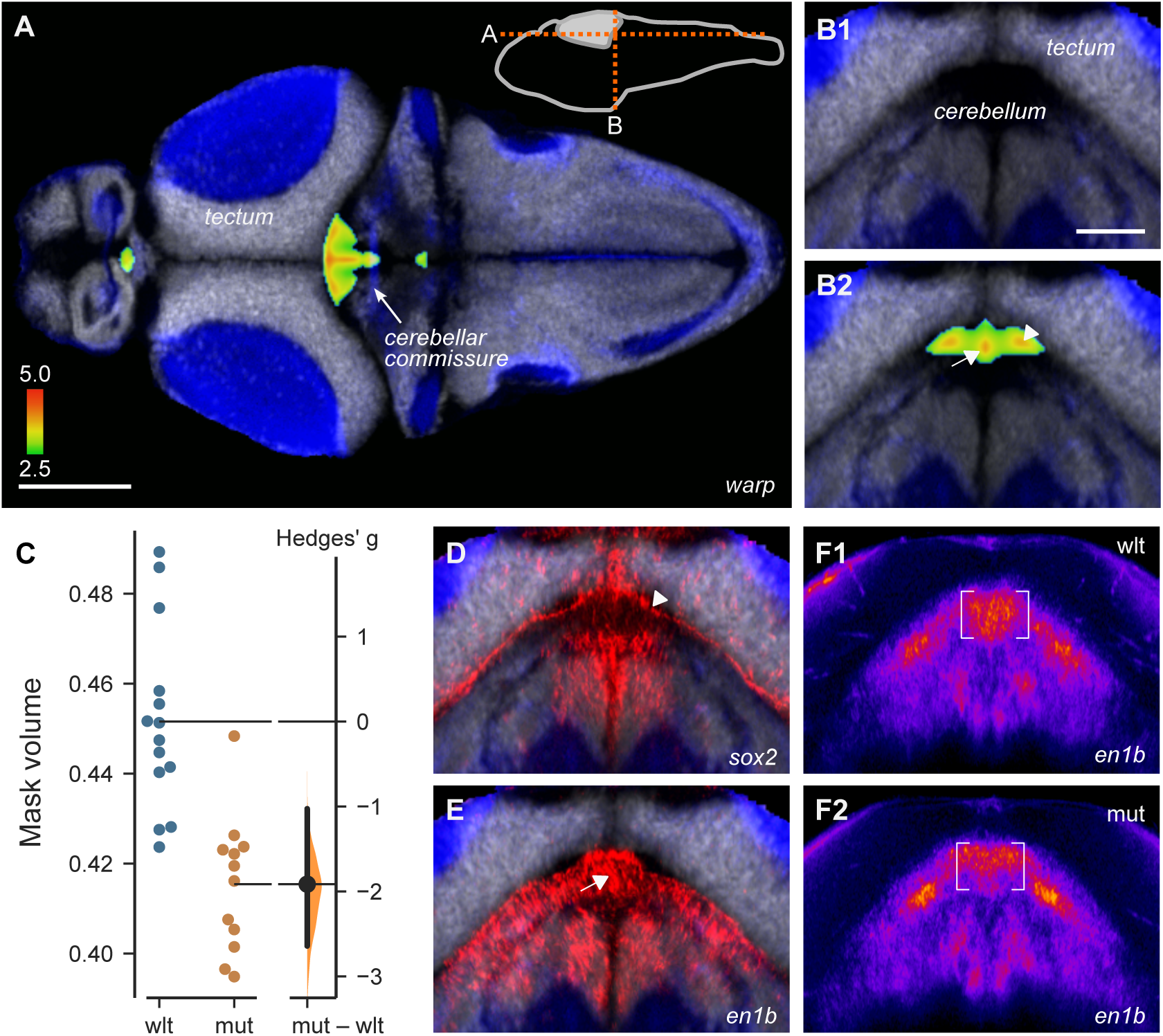
Decreased midbrain-hindbrain boundary. A-B. Voxel clusters disproportionately reduced in volume in TS2 homozygous mutants compared to wild-type siblings, shown in horizontal (A) and coronal (B) sections through the brain at the levels indicated in the inset diagram. Cellular (grey, huc:h2b-GFP) and neuropil rich (blue, huc:lyn-RFP) regions annotated using images in ZBB. Color scale shows the nominal p-value for voxels that were part of statistically significantly affected clusters. C. Volume (expressed as percent of total brain volume) of three-dimensional mask comprising the region of compressed voxels in (A), in an independent cohort of wild-type and mutant TS2 larvae. D-E. HCR fluorescent *in situ* hybridization for *sox2* (D) and *en1b* (E), at the same coronal section as in (B). Images are the mean of co-registered confocal scans from multiple larvae. Arrows in D and E are aligned with B2. F. *In situ* hybridization for *en1b* in wild-type (F1) and *y680* mutant (F2) larvae at the same coronal plane as in B. Each image is the average of 9 larvae after registration to ZBB using affine-only parameters (to prevent warping from obscuring differences). Fluorescence intensity pseudocolored to aid comparison.

In these experiments, we unexpectedly noted an increase in the *tuba:mCardinal* signal in a midline region throughout the midbrain and anterior hindbrain that normally does not contain differentiated neurons (Fig. 7A,B). To substantiate this, we designed a mask encompassing the area of increased signal in the anterior hindbrain, and measured *tuba:mCardinal* signal intensity in independent cohorts of wild-type, heterozygous and mutant larvae. Increased midline fluorescence signal was present in both mutant and heterozygous TS2 embryos compared to their wild-type siblings (Supp. Fig. 7A-B). We also performed fluorescent *in situ* hybridization for *elavl3*, a second marker of post-mitotic neurons. Mutants showed excess *elavl3* signal in the same midline region (Fig. 7C). Tissue distortion during tissue fixation and permeabilization required for *in situ* hybridization impeded quantitative replication, however a meta-analysis of three experiments confirmed that there was a significant increase in *elavl3* signal (p = 0.002; Supp. Fig. 7D). Thus, TS2 mutants inappropriately express markers associated with mature neurons in a brainstem midline region that is normally occupied by neural stem cells and neuroblasts.

**Figure 7.**
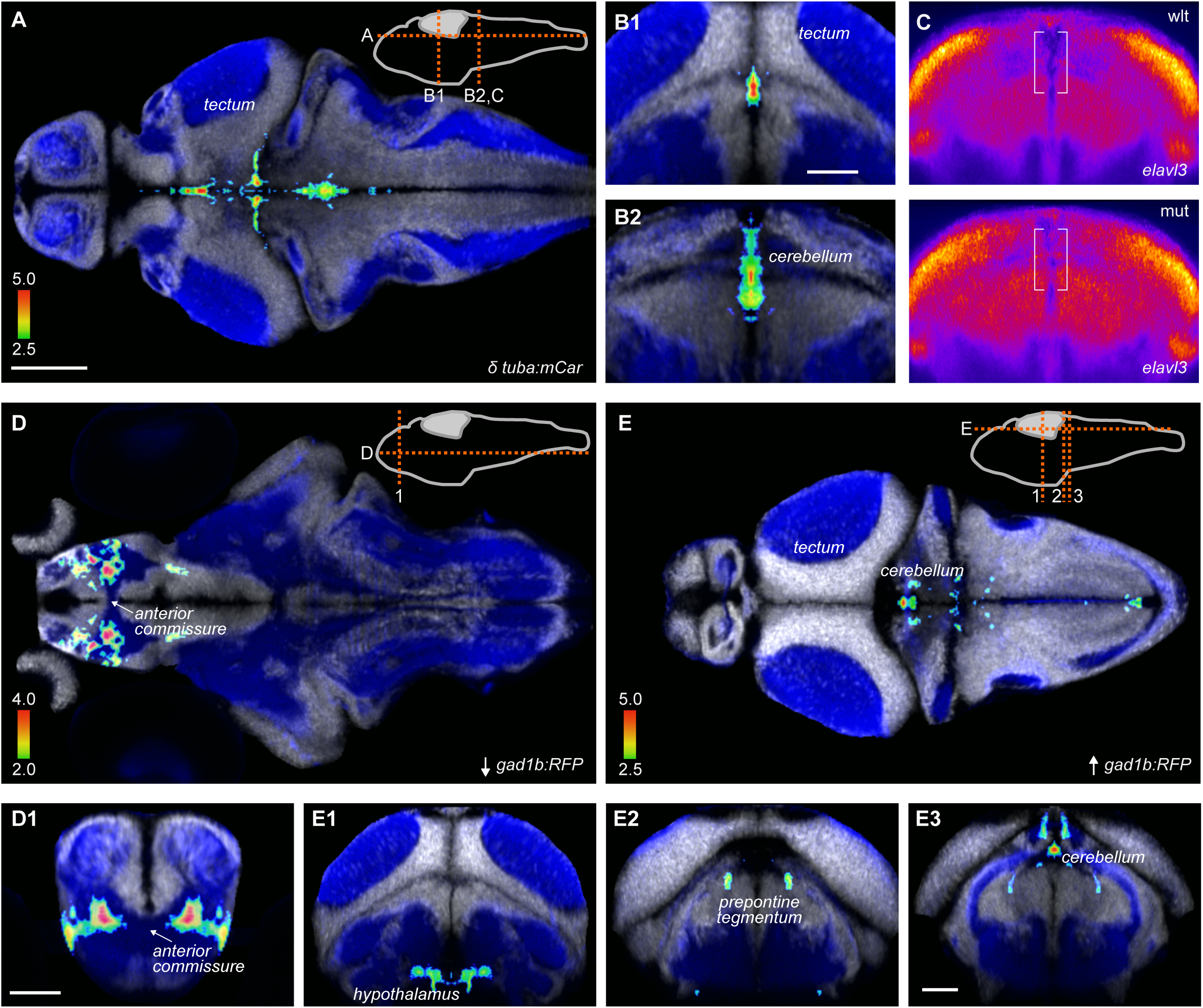
Changes in neuron distribution in TS2 homozygous larvae. A-B. Increased *tuba:mCardinal* expression in TS2 mutants, shown in horizontal (A) and coronal (B) slices, shown at the level indicated in the schematic in A. Color scale shows nominal p-value for each voxel with increased expression in mutants. Background showing cell soma rich areas (*elavl3:h2b-GFP*, white) and neuropil (*elavl3:lyn-RFP,* blue) from co-registered brain scans in ZBB, taken from MapZeBrain (*57*). C. Fluorescent *in situ* hybridization for *elavl3* in wild-type and mutant TS larvae. Each image is the mean of 9 brain scans, co-registered using affine scaling only. Coronal re-slice of confocal images shown is a 10 micron projection corresponding to the level in B2. D-E. Clusters of voxels with decreased (D) and increased (E) *gad1b:RFP* signal in TS2 mutants. Panels below show coronal re-sections at levels indicated in corresponding schematics. Scale bars: 100 um

Next, we analyzed the *gad1b:RFP* signal in mutants to examine the distribution of inhibitory neurons. Mutants showed a 5% reduction in mean fluorescence intensity in the subpallium, an effect that was also observed in a second cohort of larvae, but not in heterozygotes (Supp. Fig. 7E-G). We then combined the two cohorts by performing an ANOVA at each voxel. This disclosed that voxels with the greatest decrease in RFP signal were located in a subpallial neuropil area adjacent to the anterior commissure (Fig. 7D), and small clusters of increased *gad1b:RFP* fluorescence intensity in the cerebellum, prepontine tegmentum and hypothalamus (Fig. 7E).

### Elevated temperature reveals phenotypes in TS2 heterozygous larvae

Case reports suggest that individuals with long QT syndrome (including in one TS2 case), may be especially at risk for severe arrhythmias during febrile illness (*16, 26*). This may be because at temperatures above 37°C, Ca_V_1.2 containing L-type calcium channels open at hyperpolarized membrane potentials (*15*). We reasoned that individuals with even one TS2 gain of function allele might be highly sensitive to increased temperatures, potentially explaining why heterozygous mutations have such severe consequences in mammals, whereas heterozygous zebrafish (which are maintained at 28°C) showed relatively subtle changes. We speculated that we might reveal phenotypes in heterozygous larvae by shifting them to 37°C, a temperature that is normally well tolerated by zebrafish. Consistent with a previous report, cardiac contraction frequency in 5 dpf wild-type larvae increased to around 290 bpm at 37°C (Fig. 8A) (*27*). Atrioventricular contraction rates remained coupled and spectrograms revealed no additional frequencies. However, heterozygous and mutant larvae showed significantly elevated contraction frequencies compared to wild-types, and ventricle spectrograms revealed a range of additional frequencies suggesting arrhythmia (Fig. 8A,B; Supp Fig. 8A,B). Thus, as hypothesized, at a temperature expected to activate L-type calcium channels, heterozygous larvae exhibited a robust cardiac phenotype, whereas at lower temperatures, they resembled wild-type larvae.

**Figure 8.**
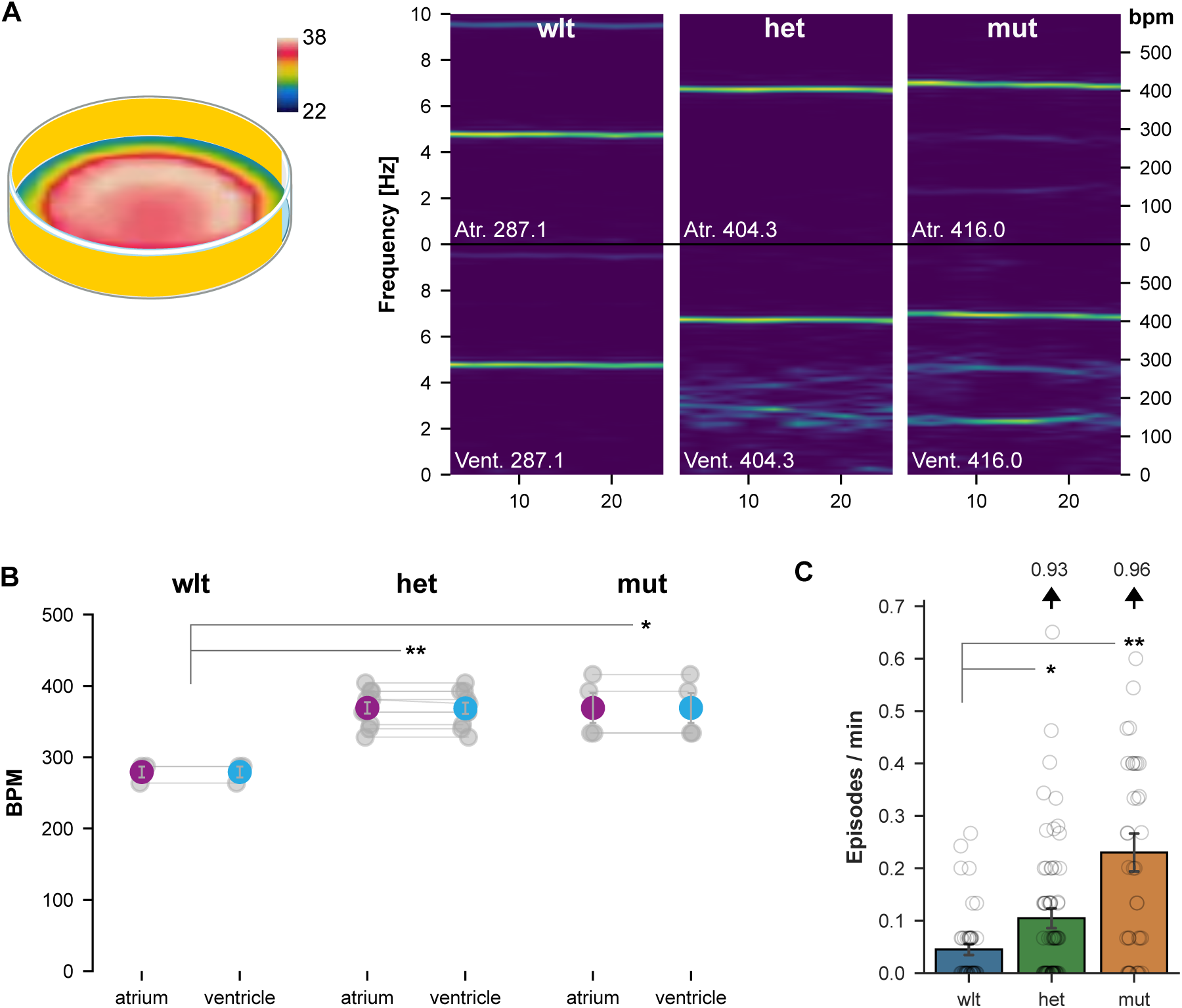
Arrhythmia and seizure-susceptibility in heterozygous larvae at elevated temperature. A. Spectrograms of cardiac contraction frequencies in wild-type, heterozygous and mutant atrium (top panels) and ventricles (bottom panels), when tested in the center of an arena warmed to 37°C (left). B. Atrium and ventricle contraction frequency for wild-type, heterozygous and mutant larvae (N = 3, 10, 4 respectively). t-test, ** p < 0.001 and * p = 0.036. C. Episode frequency of seizure-like movement for wild-type, heterozygous and mutant larvae (N = 44, 73, 37 respectively) at 37°C. Results are for three experiments combined: see Supp. Fig. 8C-D for experiments individually. Dunn’s post-hoc comparison, ** p < 0.01 and * p < 0.05.

We asked whether heat sensitivity extended to the manifestation of seizure-like activity. At 28°C, larvae that were not treated with pentylenetetrazol showed very few seizure-like locomotor episodes, irrespective of genotype (Fig. 5B, 0 mM PTZ). In contrast, at 37°C, heterozygous and homozygous mutants showed a significantly greater frequency of seizure-like episodes than wild-types, even in the absence of sensitization with pentylenetetrazol (ANOVA main effect of genotype F_2,145_ = 17.2, p < 0.001; Fig. 8C). The increase in seizure-like episodes was not part of a generalized increase in movement: during movement bouts, heterozygous and mutant embryos showed a similar locomotor speed to wild-type embryos (F_2,145_ = 0.88, p = 0.42). Elevated seizure-like behavior was greater and more robust across experiments in homozygous than heterozygous larvae (Supp Fig. 8C-D). Together, the appearance of cardiac and behavioral phenotypes in heterozygous larvae when tested at an elevated temperature suggests that normal rearing temperatures protect larvae against adverse effects of the *cacna1c* gain of function mutation and raise the possibility that increased body temperature may exacerbate symptoms in TS.

## Discussion

Here, we generated zebrafish that carry a G419R mutation in *cacna1c* as an experimental system to assess developmental and physiological effects of the gain-of-function G406R human mutation that causes Timothy syndrome 2. Cacna1c is highly conserved; in particular, the 30 amino acid residues that flank the mutation site are identical in human and zebrafish. Cacna1c^G419R^ zebrafish recapitulate key features of Timothy syndrome: bradycardia, episodic arrhythmia and seizure susceptibility. Homozygous mutant TS2 zebrafish showed a reduction in the midbrain-hindbrain boundary and, correspondingly, a reduction in the size of adjacent midbrain and hindbrain areas. Several neuropeptide genes showed elevated expression in mutants, including *bdnf* and *vgf*, genes implicated in the control of neurogenesis. Although TS is a dominant disorder, heterozygous zebrafish showed only subtle abnormalities in brain morphology and transcription but normal survival and fertility. Under standard conditions, heterozygotes did not show arrhythmia or seizure susceptibility, but these phenotypes emerged when body temperature was raised to 37°C, a point at which Ca_V_1.2 channels open spontaneously.

Signature features of TS include a prolonged QT interval, syndactyly and neurodevelopmental abnormalities including autism and seizures (*19*). TS2, which arises from a G406R mutation in exon 8 rather than exon 8A, is clinically similar but without syndactyly. To assess the extent to which deficits in Cacna1c^G419R^ zebrafish resemble clinical manifestations in TS2, we focused on cardiac and seizure phenotypes, which have been extensively characterized in other zebrafish models. TS2 mutants exhibited bradycardia in 2 dpf embryos and intermittent arrhythmia similar to 2:1 atrioventricular block (AVB) in older larvae. Bradycardia and AVB are present in a majority of TS2 patients perinatally (*14*). Behavioral deficits in TS patients include autism, seizures and – in at least a few reports – auditory sensitivity. To test seizure susceptibility, we exposed larvae to a low dose of the GABA receptor antagonist pentylenetetrazol. Homozygous (but not heterozygous) TS2 larvae treated with pentylenetetrazol showed an increase in the number of seizure-like locomotor events compared to siblings. Together, the presence of specific cardiac arrhythmias – bradycardia and 2:1 AVB – and seizure susceptibility indicate that TS2 larvae recapitulate key TS clinical manifestations and may therefore provide a useful tool for investigating how gain-of-function mutations in this disorder disrupt normal development.

During neural development, there is a switch from the inclusion of exon 8A in neural progenitors to the inclusion of exon 8 in mature neurons (*2, 3*). It was recently shown that the use of an antisense inhibitory oligonucleotide to prevent the inclusion of exon 8A increased exon 8 inclusion and rescued neurodevelopmental phenotypes in a TS1 brain organoid model (*8*). Injection of morpholino antisense oligonucleotides into zebrafish embryos is routinely used to block pre-mRNA splicing (*28*), but it was not clear whether a similar strategy would be effective in zebrafish TS2 mutants where the mutation affects exon 8, which shows more than 10-fold greater utilization than exon 8A throughout development. However, a morpholino against the intron 7 - exon 8 boundary was highly effective in switching isoform expression from exon 8 to exon 8A. Cardiac contraction frequency in exon 8A-containing larvae was increased irrespective of genotype, such that mutants injected with the splice-blocking morpholino showed similar contraction frequency to wild-types injected with a control morpholino. This supports the idea that splice-modifying antisense oligonucleotides might offer a therapeutic approach in Timothy syndrome.

Consistent with a key role for intracellular calcium signaling in regulating gene expression in neurons (reviewed in (*29, 30*)), gene expression in neurons derived from Timothy syndrome patient stem cells and stem cells from healthy controls differed considerably (*31*). Accordingly, RNA-seq data from TS2 mutants showed widespread changes, prominently in the expression of genes that encode neuropeptides. A known downstream target of calcium influx through *cacna1c* containing L-type calcium channels is *bdnf* (*32*). RNA-seq revealed elevated expression of several neuropeptides in mutants, including *bdnf*, its downstream factor *vgf* and *si:dkey-175g6.2,* a second homolog of *vgf* present in zebrafish, which was also elevated in TS2 heterozygous larvae. Increased expression of these neuropeptides may contribute to changes in neurogenesis. We also observed decreased expression of *gcgb* (*glucagon b*) in mutants, consistent with reports of lower glucagon immunoreactivity and impaired glucagon secretion during hypoglycemia in a TS mouse model and possibly contributing to life-threatening hypoglycemia episodes in patients (*9*). Thus, *cacna1c* may have a conserved role in regulating energy metabolism in vertebrates.

We found that *cacna1c* is broadly expressed throughout stages of neural development in zebrafish, with conspicuous expression in the cerebellum. The relative expression levels of *cacna1c* over time paralleled that of the mature neuron marker, *sypa*; however, other evidence suggests that *cacna1c* may also regulate early developmental processes. Brain size was diminished in mutants, and in particular the total volume of brain regions that are predominantly cellular was reduced compared to the total volume of neuropil-rich regions. There was a disproportionately large reduction at the midbrain-hindbrain boundary (MHB) and a matching loss of *en1b* expression, a MHB marker. The MHB controls the development of the adjacent cerebellum and optic tectum, and consistent with this, we saw a significant reduction in cell density in the major neuron-containing area of the optic tectum, the stratum periventriculare. In parallel, we observed an increase in the expression of two mature neuron markers – the transgenic line *tuba:mCardinal*, and *in situ* hybridization for *elavl3* – adjacent to the brain midline in mutants, a region that normally includes stem cells and proliferating neuroblasts. L-type calcium channels have been implicated in regulating neuronal proliferation (reviewed in (*33*)). In humans, *CACNA1C* is strongly expressed in the outer subventricular zone, where it may contribute to the depolarizing effects of GABA on neural progenitor cells and regulate cell fate decisions (*2*). Premature terminal differentiation of proliferating neuroblasts might explain the smaller overall brain volume in mutants, with increased neuron signal at the midline. Indeed, *bdnf* and *vgf*, genes that were upregulated in TS2 mutants, are known to regulate neurogenesis (*21, 22*). However, LTCCs are also known to regulate neuronal migration (*34*). TS2 mice showed an increase in tangential migration of inhibitory neurons from the median ganglionic eminence (MGE) (*5*). Conversely, electroporation of *Cacna1c* mRNA with the TS mutation suppressed the radial migration of excitatory neurons into the upper cortical layers in mice (*35*), and in an organoid model, neurons from TS1 patient-derived stem cells showed delayed migration of MGE interneurons (*6, 7*). On the other hand, neither inhibition nor activation of L-type calcium channels affected the migration of newborn neurons in the olfactory bulb of postnatal mice (*36*). The effects of LTCC activity on neuronal migration appear to vary with stage and cell type in the developing brain, and thus, the expression of mature neuron markers at the midline region in TS2 mutants might reflect premature terminal differentiation of neuroblasts into neurons, suppressed neuronal migration, or both.

An unexpected limitation of the zebrafish model was that significant cardiac and behavioral changes were detected only in homozygous mutants, whereas patient mutations are dominant. A further discrepancy is that TS2 larvae could be raised with only minor loss of viability to fertile adults, compared to high mortality in patients. It is not uncommon for mutations that cause haploinsufficient psychiatric disorders to be homozygous viable in zebrafish (*37*), however, given the severe effects in TS patients we considered whether phenotypic effects of the mutation were compensated or masked in zebrafish. It is unlikely that a redundant gene mitigates *cacna1c* mutations because mutations that disrupt *cacna1c* are lethal in zebrafish due to a severe impairment in the production of ventricular cardiomyocytes (*12*). Patient mutations affect the second to last nucleotide of exon 8 or 8A, and in an induced pluripotent stem cell model, this disrupts splicing leading to a shift in isoform use (*2*). However, we did not detect differences in isoform expression or splice site selection in TS2 mutants. Of the 8 other L-type voltage-gated calcium channel genes in zebrafish, the only significant change at either 2 or 6 dpf was a 10% increase in expression of *cacna1sb* at 2 dpf, making it unlikely that genetic compensation explains the survival of TS2 mutants. We therefore think that cellular adaptation to the mutation is unlikely to explain why heterozygous zebrafish show only mild phenotypes.

Instead, we speculate that biological and/or environmental factors may account for heterozygous viability. First, in the human heart, L-type calcium channels constitute more than 90% of all calcium channel transcripts, compared to only 35% in zebrafish, where the majority are T-type calcium channels (*38*). Second, in human cardiomyocytes, calcium entering through channels in the cell membrane triggers calcium release from the sarcoplasmic reticulum, potentially amplifying effects of changes in extracellular calcium influx. In contrast, zebrafish ventricular myocytes primarily rely on extracellular calcium influx for contraction (*39*). Third, it has also been argued that due to intrinsic differences in the cardiac conduction system and repolarization properties in fish, delayed repolarization of cardiomyocytes is less likely to cause arrhythmia than in mammals (*40*). These factors may mean that zebrafish are simply less sensitive to activating mutations in *cacna1c*. However, we directly tested another idea – that zebrafish are protected from deleterious effects by normal 28°C rearing conditions. At temperatures greater than 37°C, Ca_V_1.2 channels are active at hyperpolarizing potentials, driving intrinsic neuronal firing (*15*). TS mutant mammalian cells may therefore be especially prone to sustained depolarization. To test this idea, we measured cardiac function in TS2 larvae held at 37°C. In contrast to measurements at 28°C where a phenotype occurred only in homozygous mutant larvae, at 37°C both mutant and heterozygous larvae showed atrial and ventricular tachycardia. As TS patients are prone to infections, elevated L-type calcium channel activity during fever may lead to tachyarrhythmia, which is a major cause of mortality, and in fact infection has been reported as an associated factor for sudden death in TS1 (*14*). Our data suggest the importance of studying whether patient outcomes may be improved by close monitoring and management of body temperature.

## Methods

### Husbandry

Embryos and larvae were raised in a 14:10 light:dark cycle at 28°C (except as otherwise noted) in E3 medium supplemented with 1.5 mM HEPES pH 7.3 for buffering. Transgenic lines used in this study were *TgBAC(gad1b:lox-RFP-lox-GFP)nns26* (*gad1b:RFP*) (*41*) and *Tg(Cau.Tuba1a:MCardinal)y516* (*tuba:mCardinal*) (*11*). Experiments were approved by the NICHD Animal Care and Use Committee.

### CRISPR-Cas9 mutagenesis and fish line maintenance

A four-nucleotide substitute that leads to a G419R amino acid substitution in the zebrafish CACNA1C protein was introduced into the genome of ABC strain fish by CRISPR-Cas9 knockin (In Vivo Biosystems, Eugene, OR). This allele, NM_131900.1(cacna1c):c.1252_1255delAGCGinsTCTA, designated *y680*, was originally made in an ABC background, and maintained by repeatedly crossing heterozygous carriers to TL wild-type stock (Zebrafish International Resource Center, Eugene, OR). For genotyping, we performed PCR with DNA samples from tail fins and primers 5-GTTTCCGCTCACTGTCTTCC and 5-ACACGCATTTAGCACACAATGCTTAtC, then digested PCR products with XbaI for 2 h at 37°C and separated on a 1% agarose (biotechnology grade, VWR) + 2% MetaPhor agarose (Lonza) gel. The wild-type 202-bp PCR product was intact, while mutant PCR products were cleaved to a 170-bp fragment.

### Survival monitoring

Fish were genotyped as larvae, then raised with siblings of the same genotype in groups of 4–6 per 1.8 L tank and counted daily. For the survival monitoring in large groups, fish were genotyped at the larval stage and placed into 1.8 L tanks in groups of 20–29 (2 groups of 10 heterozygotes, 5 wild-type, 5 mutants, and 1 group of 12 heterozygotes, 7 mutants, 10 wild-type) at the age of 6 dpf. Fish were counted and genotyped after they reached the adult stage. Photographs of the adult fish taken from above were processed in ImageJ to measure body length (from the tip of the mouth to the base of the caudal fin).

### Expression analysis

To compare expression of *cacna1c* to markers of stages of neuronal maturation, we downloaded the DanioCell single cell sequencing dataset from NCBI GEO (accession GSE223922, (*17*)) and extracted cells annotated as ‘neural’. For each gene of interest, we normalized expression to total UMIs per cell, then calculated mean expression across cells of a given age range. Expression values for each gene were normalized to maximal expression across all ages then smoothed for plotting.

### RNA analysis

To prepare samples for transcriptomic analysis, embryos (2 dpf) and larvae (6 dpf) were cut in the middle of the yolk extension under terminal anesthesia. Tail tissue was used for DNA isolation and genotyping, and the anterior part of the body (mostly head) stored in RNAlater (ThermoFisher) at –20°C. Three heads from larvae of the same genotype were pooled, and RNA was isolated in a Maxwell 16 instrument with a Maxwell 16 LEV Simply RNA Tissue kit (Promega). Capillary electrophoresis in an Agilent Bioanalyzer was used for RNA quality control. cDNA libraries were prepared and sequenced at the Molecular Genomics Core, NICHD. We used FastQC v.0.12.1 to check FASTQ file quality and trimmed sequences with Cutadapt v.4.4 before aligning to zebrafish genome version GRCz11 using STAR v2.7. We then used featureCounts from the Subread package v2.0.3 to generate gene counts from BAM files. Genetic background associated polymorphisms that control gene expression levels frequently cause differences in expression of genes on the same chromosome as an experimental mutation that are linked to the mutation rather than being effects of the mutation itself (*42*). We expected this phenomenon to be exacerbated in TS2 larvae because the *y680* mutation was made on an ABC wild-type background, and subsequently crossed to a TL background. Indeed, differentially expressed genes (DEGs) were disproportionately located on chromosome 4, the same chromosome as *cacna1c* (70.6%, Supp. Fig. 3A). Therefore, for annotation of DEGs and Gene Ontology pathway analysis, we excluded genes located on chromosome 4. We used DESeq2 v1.33.0 with apeglm shrinkage to measure differential expression. We removed genes with less than 100 mean counts, and applied the Benjamini-Hochberg false discovery rate (FDR) correction to DESeq2 p-values and considered genes with an FDR of < 0.1 as significant. Gene pattern analysis was performed with DEGreport version 1.34.0, utilizing the ConsensusClusterPlus algorithm to cluster genes based on similar expression profiles, using differentially expressed genes between wild-type and mutant as input (FDR < 0.1, but not filtering for base mean expression) (*43*). For gene ontology analysis, lists of ENSEMBL identifiers for upregulated and downregulated genes were processed with the ShinyGO web tool (https://bioinformatics.sdstate.edu/go/; (*44*)) with FDR cutoff set to 0.05, number of pathways to show set to 500, and “remove redundancy” option flagged. Daniocell (daniocell.nichd.nih.gov) and zfin.org databases were used to annotate the functions and tissue specificity of expression of the differentially regulated genes (*17, 45*). To compare splicing in wild-type and mutant larvae, we used *featureCounts* from the subread package to measure splice junctions, then normalized the junction counts for each sample using scaling factors generated by deepTools. Independent RNA samples prepared in the same manner were used for quantitative RT-PCR to characterize the usage of mutually exclusive *cacna1c* exons 8 and 8A at 2, 4, and 6 dpf. cDNA synthesis and qPCR were performed at BioInnovatise (Rockville, MD). Primers are listed in Supplementary Table 1, including for the reference gene *eef1a1l1* (*EF1α*) for which we used previously described primers (*46*).

### Heart rate measurements

Larval fish were anesthetized with tricaine to prevent spontaneous movements and placed under a microscope connected to a Chronos 1.4 high-speed camera, allowing heart videos to be recorded at 100 frames per second. We used ImageJ to calculate mean pixel intensity over time in ROIs that covered the atrial/ventricular wall (*47*). Time series data was processed in Python to compute spectrograms (which represent the power present at different frequencies in the Fourier transform of mean image intensity from each cardiac chamber over time) using the *spectrogram* function in the scipy.signal library (*48*). We used the *find_peaks* function in scipy.signal to determine mean ventricular and atrial contraction rates for experiments at 28°C but found this was not sufficiently accurate at high contraction frequencies for larvae recorded at 37°C, so instead, we extracted the dominant contraction frequency using the Fourier transform in scipy.signal *spectrogram*. To measure contraction frequency at 37°C we placed larvae in the center of a glass Petri dish wrapped with a foil Kapton flexible heating element (Omegalux) powered by a TC200 Temperature Controller (Thorlabs), with a 10 kΩ thermistor to measure temperature. We verified the temperature in the center of the dish using an I3 infrared thermal imager (FLIR). Exposure to 37°C is widely used to activate transgenic heat-shock promoters in zebrafish larvae and does not cause negative effects (*49*).

### Morpholino injections

To assess morpholino efficacy, fertilized wild-type embryos were injected with standard control morpholino (Gene Tools) or 1 ng or 3 ng of cacna1c-i7e8 morpholino (5-CATCCTGCATCTGAGACGAGGAGGG). At 2 and 5 dpf, embryos were dissected caudal to the yolk, and anterior tissue (including head and heart) was collected for mRNA preparation using the Maxwell RSC simplyRNA kit. We used SuperScript™ III One-Step RT-PCR System with Platinum™ Taq High Fidelity DNA Polymerase with primers 5-TTTGCCATGCTGACTGTGTT and 5-GTGATCCAGTCCAGGTAGCC producing a 275 bp band for either exon 8 or exon 8A containing isoforms. We then digested the product with EcoRI: exon 8 has no EcoRI target, whereas exon 8A is cleaved into 195 and 80 bp fragments. A dose-dependent shift in exon 8 splicing was apparent at 2 dpf (Supp. Fig. 4B) but not at 5 dpf (not shown). For testing cardiac contraction frequency, wild-type and mutant cousin embryos (ie, embryos that were derived from parents that were wild-type and mutant siblings respectively) were injected with 3 ng control or i7e8 morpholino, then tested at 2 dpf.

### Brain morphometry

For brain morphometry experiments *cacna1c*^y680+/-^*; gad1b:RFP; tuba:mCardinal* fish were backcrossed to *cacna1c*^y680+/-^, and fin clips from offspring were collected at 3 dpf for genotyping. Brains of live larvae were imaged at 6 dpf on a Nikon A1 confocal microscope with a 20× water immersion objective, voxel size 1×1×2 μm. Z intensity correction was used to compensate for weaker signals from the ventral-most parts of the brain. Using ImageJ, confocal stacks were rotated to orient the main axis of the brain horizontally and separate channels into individual files. Stacks were then aligned to the *gad1b:RFP* and *tuba:mCardinal* patterns from ZBB using multi-channel registration in ANTS (*50*). After registration, we calculated the log Jacobian determinant of the transformation matrix to generate a map for each fish, showing the extent to which individual voxels were contracted or dilated to match the reference brain (LJD channel). Next we used Cobraz to identify clusters of voxels that differed in wild-type and mutant larvae in the *gad1b:RFP*, *tuba:mCardinal* or LJD channels, by using permutation analysis to rigorously establish significance thresholds for each channel (*11*). We repeated this procedure in an independent cohort of larvae and only reported changes that were consistent between the experiments. In addition, we symmetrized maps of significant voxels by taking the greater p-value at each bilaterally symmetric position. To measure total brain volume, neuroanatomical annotations in the Zebrafish Brain Browser reference brain were inverse-transformed onto the original brain scans. Similarly, to measure changes in cerebellar or midline volume in mutants, we created masks covering the affected region in the first experiment and, in the second cohort of fish, inverse-transformed it back onto the unprocessed brain scans, allowing us to count voxels within the mask in wild-type and mutant brains and thereby determine the volume of the affected area. Imaging and brain morphometry procedures were as previously described (*51*).

### Brain activity measurements

To measure neural activity, we first crossed heterozygous TS2 larvae to *Tg(elavl3:H2B-GCaMP6s)jf5* (*52*), then backcrossed to TS2 heterozygotes. We embedded larvae in agarose and used epifluorescent imaging to record neural activity at 0.5 Hz for 5 min. We then extracted DNA for posthoc genotyping. To correct for drift over the recording period, we used ImageJ to perform a rigid registration on slices in the virtual stack. We then used ANTs to co-register recordings. We applied a 2 sigma Gaussian filter from *scipy.ndimage* to correct for subpixel drift and noise in each image. To account for differences in GCaMP expression between fish, we used the minimum fluorescent signal at each voxel over the 5 min recording period as its baseline, found the largest change in fluorescence at each voxel as the maximum magnitude change, and calculated the percent change over baseline (dF/F). We used the mean dF/F within a mask that encompassed cellular brain regions as a measure of spontaneous neural activity.

### Hybridization chain reaction

(HCR) fluorescent *in situ* hybridization to detect gene expression was performed according to the manufacturer’s protocol (Molecular Instruments) with the proteinase K treatment step omitted and the samples partially dissected (tail cut off just rostrally to the yolk extension under terminal anesthesia prior to fixation). Oligonucleotide probes for *cacna1c, vgf, sox2* and *en1b* were used with the B3 amplifier, and probes for *elavl3* were used with B1 or B2 amplifier. Probes for *elavl3* with the B2 amplifier were from Molecular Instruments, and other probes were designed in-house (*53*). Samples were stored in PBS pH 7.4 at 4°C prior to imaging. Probe sequences are in Supplementary File 1.

### Behavioral analysis

To measure seizure-like activity, larvae derived from a heterozygous incross were transferred to individual wells of a 36-well testing arena filled with E3 embryo medium with 1% DMSO (vehicle control) or freshly prepared 1 mM PTZ in E3 with 1% DMSO. Wells had dimensions 7 × 7 mm. After 30 minutes exposure to PTZ, locomotor activity was then recorded for 15 minutes; then the light was extinguished and activity recorded for another 15 minutes. Larvae were tracked in real-time at 10 Hz using DAQtimer then post-hoc genotyped (*54*). After recordings were complete, we calculated the total distance moved for each larva in 1 second windows, then calculated a large-movement distance threshold that was exceeded in only 0.1% of windows in wild-type vehicle-treated larvae – thus over a 15 minute recording period, we would expect slightly less than one such movement in untreated wild-type larvae. We then recorded the total number of instances in which this threshold was exceeded in 1 second windows for each larva in illuminated and dark conditions. To assess startle behavior and prepulse inhibition, the responses of the larvae to vibrational stimuli were filmed at 1000 frames per second with a high-speed camera and analyzed with Flote software (*55*). We performed two replications of this experiment, and for each measure we assessed significance using ANOVA to calculate the main effect of genotype, controlling for the experiment and controlling the false discovery rate with Benjamini-Hochberg correction. To assess activity for larvae at 37°C, we used the same temperature controller as for cardiac contraction experiments, with larvae in a 4-well grid inside the chamber, with each well having the same dimensions as above.

### Statistical analysis

Data were analyzed using in Python 3.7 and R (www.R-project.org). Estimation plots were generated using the *dabest* package and represent means and effect size (Hedges’s g) (*56*). All *t* tests are two-tailed and independent samples.

## Acknowledgments

This work was supported by the Intramural Research Program of the *Eunice Kennedy Shriver* National Institute for Child Health and Human Development (NICHD) and utilized the high-performance computational capabilities of the Biowulf cluster at the National Institutes of Health, Bethesda, MD.

## Author contributions

S.A.S. - investigation, formal analysis, writing – original draft, writing – review and editing

A.S. - investigation, formal analysis J.S., K.R., K.L., S.K. - investigation

D.N. - formal analysis, software, visualization M.M., G.M. - formal analysis, software

A.G. - conceptualization, funding acquisition

B.F. - conceptualization, investigation

H.A.B. - conceptualization, investigation, funding acquisition, software, supervision, visualization, writing – original draft, writing – review and editing

## Competing interests

Authors do not have any competing interests

---

*Supplementary video 1. Elevated brain activity in mutants*

Fluctuating GCaMP fluorescence signal in example wild-type (top) and mutant (bottom) TS2 larvae recorded over 5 min.

*Supplementary video 2. Atrium and ventricular contractions*

Videos showing atrial (blue outline) and ventricular (orange outline) contractions in wild-type (A) and mutant (B) TS2 larvae, slowed to half speed.

*Supplementary File 1. Oligonucleotide sequences for HCR probes*

Oligonucleotide sequences for probes used to perform HCR fluorescent *in situ* hybridization.

**Supplementary figure 1.**
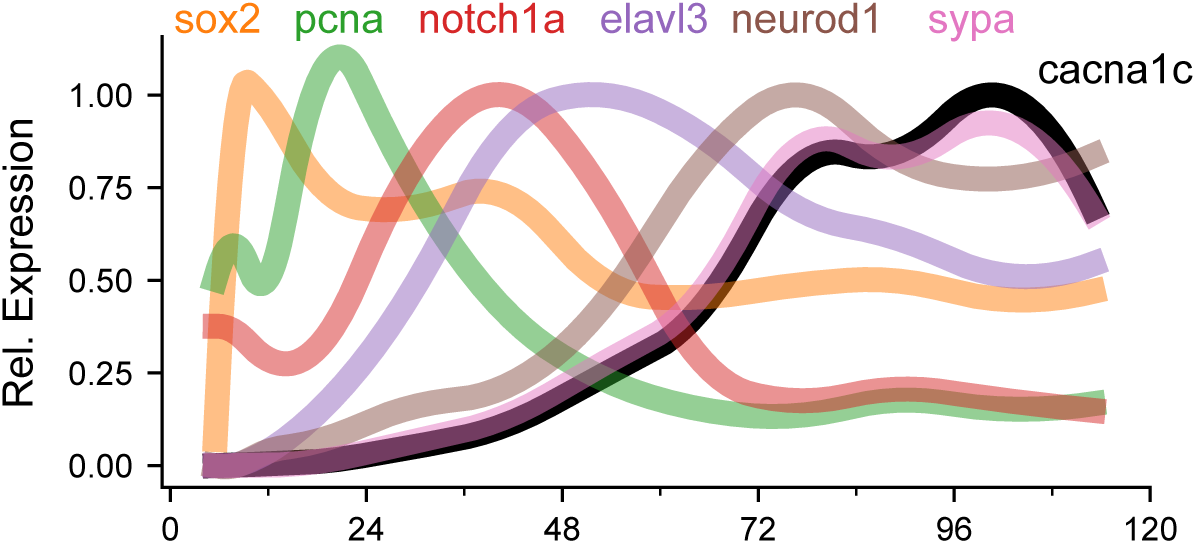
Cacna1c expression during brain development in zebrafish. Expression of *cacna1c* during embryonic and early larval brain development (0-120 hpf) compared to markers of neuronal maturation. Data is mean read counts per stage, using cells annotated as neural in DanioCell, smoothed and normalized to maximum expression for each gene.

**Supplementary figure 2.**
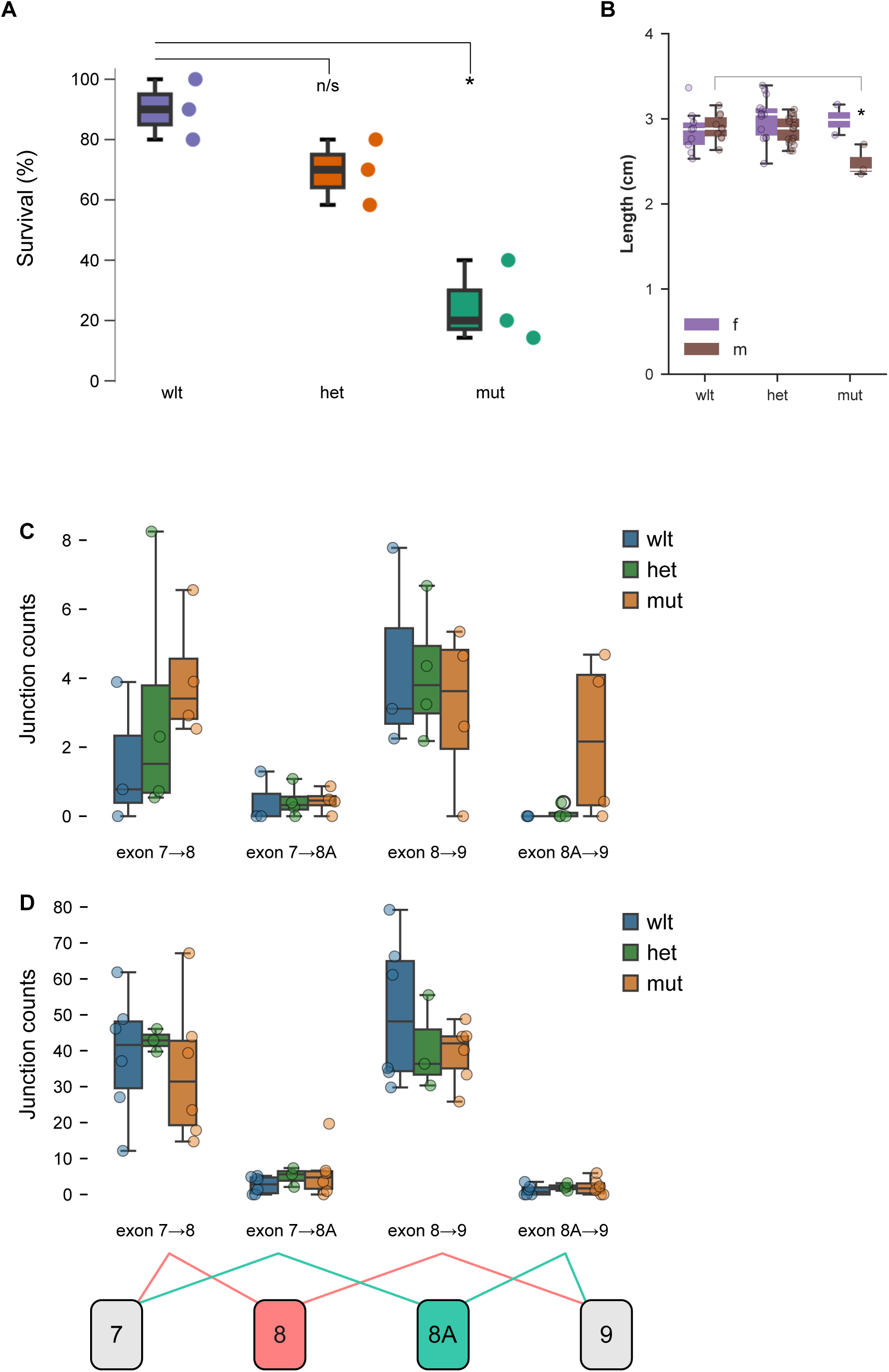
Characterization of TS2 zebrafish. A. Survival by genotype for sibling larvae raised in mixed groups of 20–30 individuals. N = 3 groups per genotype. * Tukey’s test, p = 0.001. B. Length of adult fish by genotype and sex (m, male; f, female). * t-test, p < 0.05. C-D. Normalized junction counts from RNA-seq for exon 7 to either exon 8 or 8A, and exon 8 or 8A to exon 9 in wild-type, heterozygous and mutant TS2 larvae, at 2 dpf (C) and 6 dpf (D). Between-group differences not significant for any junction.

**Supplementary figure 3.**
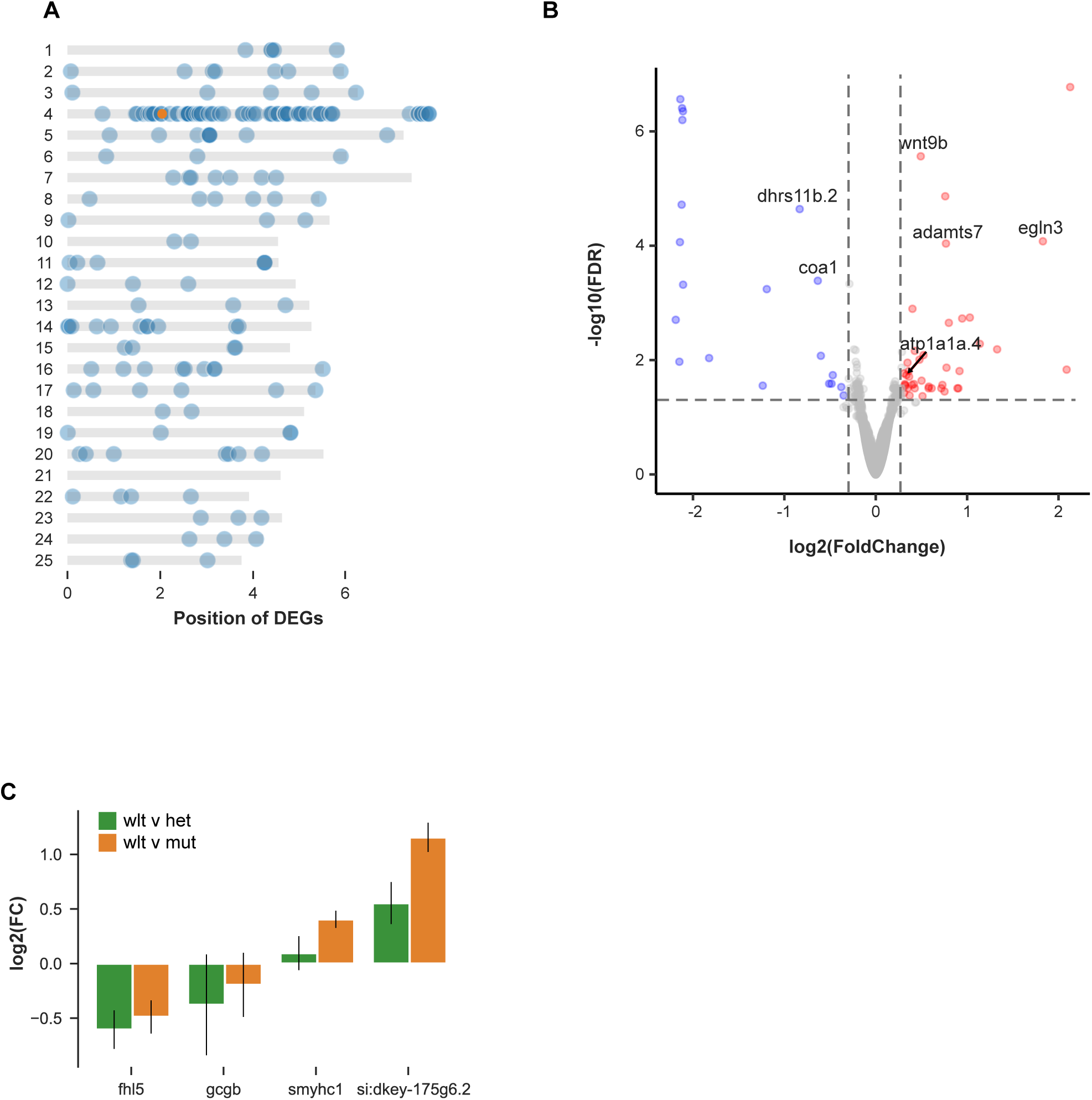
Differentially expressed genes in TS2 mutants. A. Chromosomal location of differentially expressed genes in TS2 mutant larvae compared to wild-type siblings (blue markers). Location of *cacna1c* indicated (orange marker). B. Volcano plot for genes differentially expressed between wild-type and mutant TS2 in 2 dpf larvae using RNA-seq of anterior tissue section (head and heart). C. Change in expression for four genes that were significantly different between heterozygous and wild-type siblings larvae, that were also nominally different in the wild-type/homozygote comparison.

**Supplementary figure 4.**
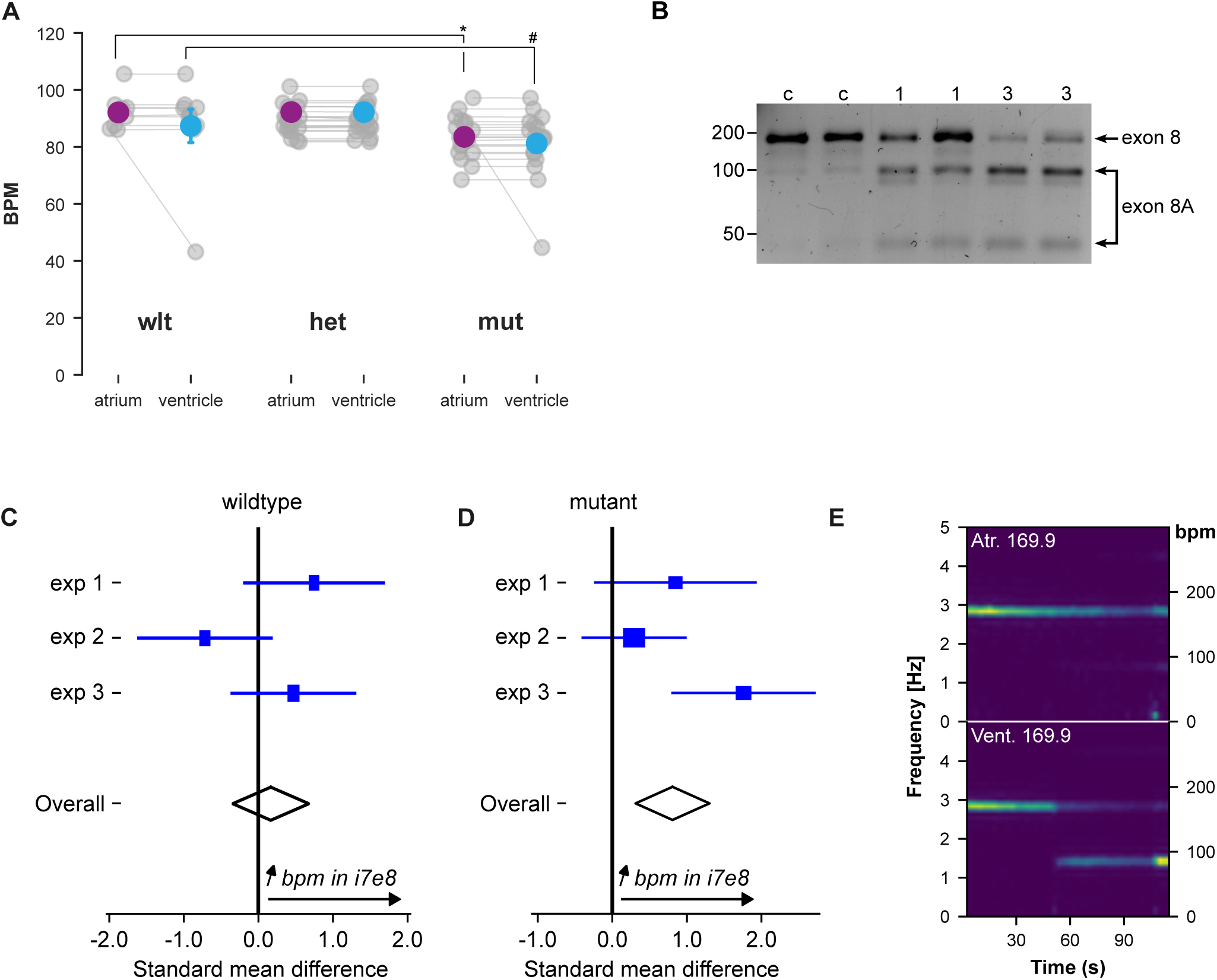
Ventricular bradycardia and antisense control of exon 8 splicing. A. Contraction frequency (beats per minute, bpm) in sibling wild-type, heterozygous and homozygous mutant larvae. Atrial frequency was significantly reduced in mutants (*, t-test p = 0.0036) but not in heterozygous larvae. For ventricle frequency, because of outlier individuals, we used a Mann-Whitney test to compare: #, p = 0.016. B. RT-PCR and restriction digest for 2 dpf embryos injected with control morpholino (c), 1 ng per embryo i7e8 morpholino (1) or 3 ng i7e8 morpholino (3). C-D. Meta-analysis of three experiments injecting morpholinos into wild-type (C) and mutant (D) embryos. Standard mean difference and confidence interval for atrial contraction frequency between control and i7e8 morphants is shown for each experiment, with diamonds showing overall effect, which was significant in mutants (standard mean difference 0.81, confidence interval [0.31, 1.3]). E. Spectrogram for an individual mutant larva at 6 dpf, where the atrium:ventricle contraction frequency shifted from 1:1 to 2:1 during a two minute recording.

**Supplementary figure 5.**
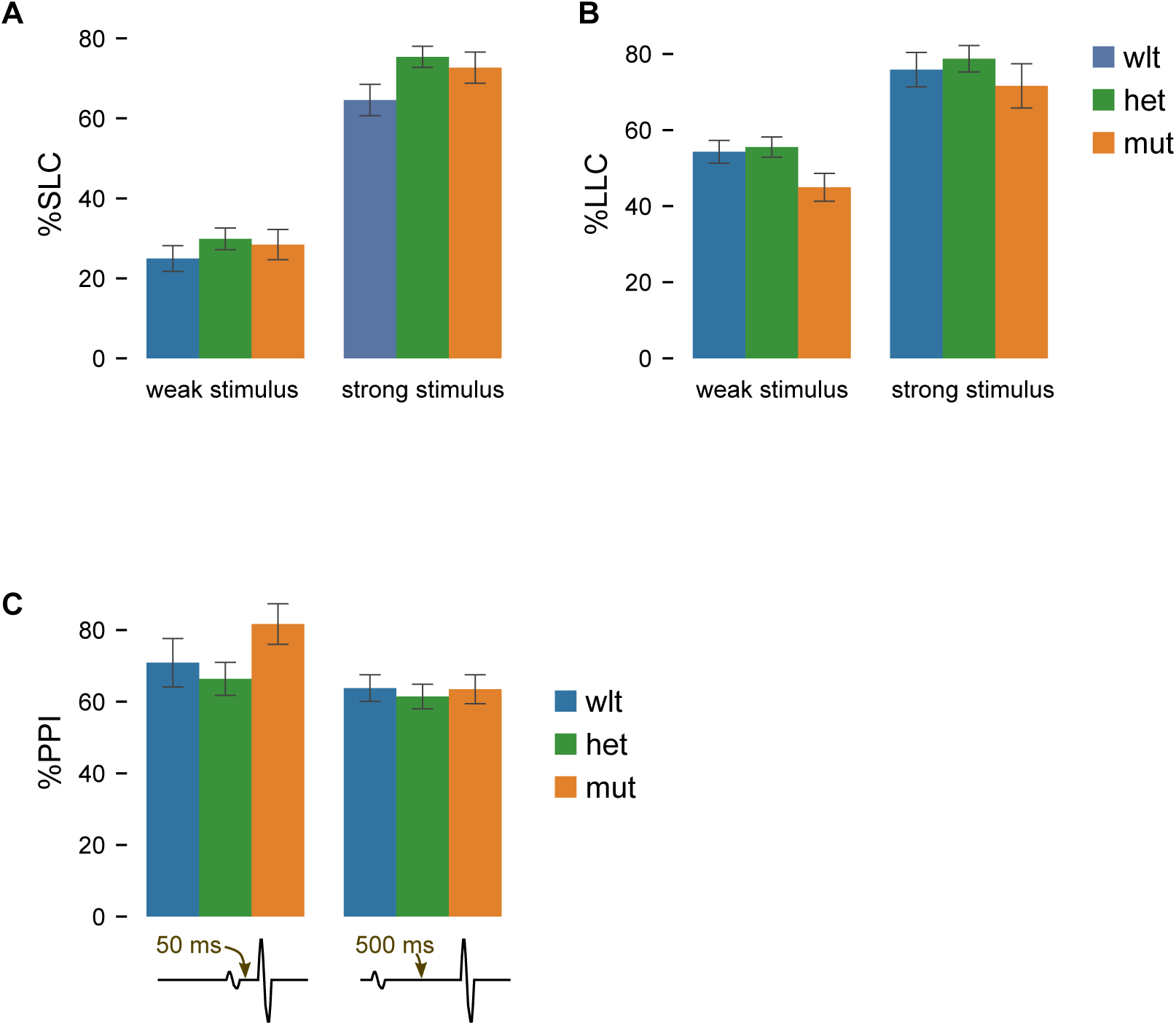
Auditory sensorimotor processing. A-B. Auditory C-start responsiveness (A) and prepulse inhibition (B) in *cacna1c* wild-type, heterozygote and mutant larvae (N= 48, 82 and 37). Larvae perform two distinct types of auditory C-start response distinguished by latency: short latency C-starts (left) and long-latency C-starts (right). No significant effects of mutation were observed for either mode of response, either at low or high intensity auditory stimuli. C. Prepulse inhibition of the startle response at 50 ms (left) and 500 ms (right) interstimulus intervals between prepulse and startle inducing stimulus, tested in the same larvae as above. No differences are significant.

**Supplementary figure 6.**
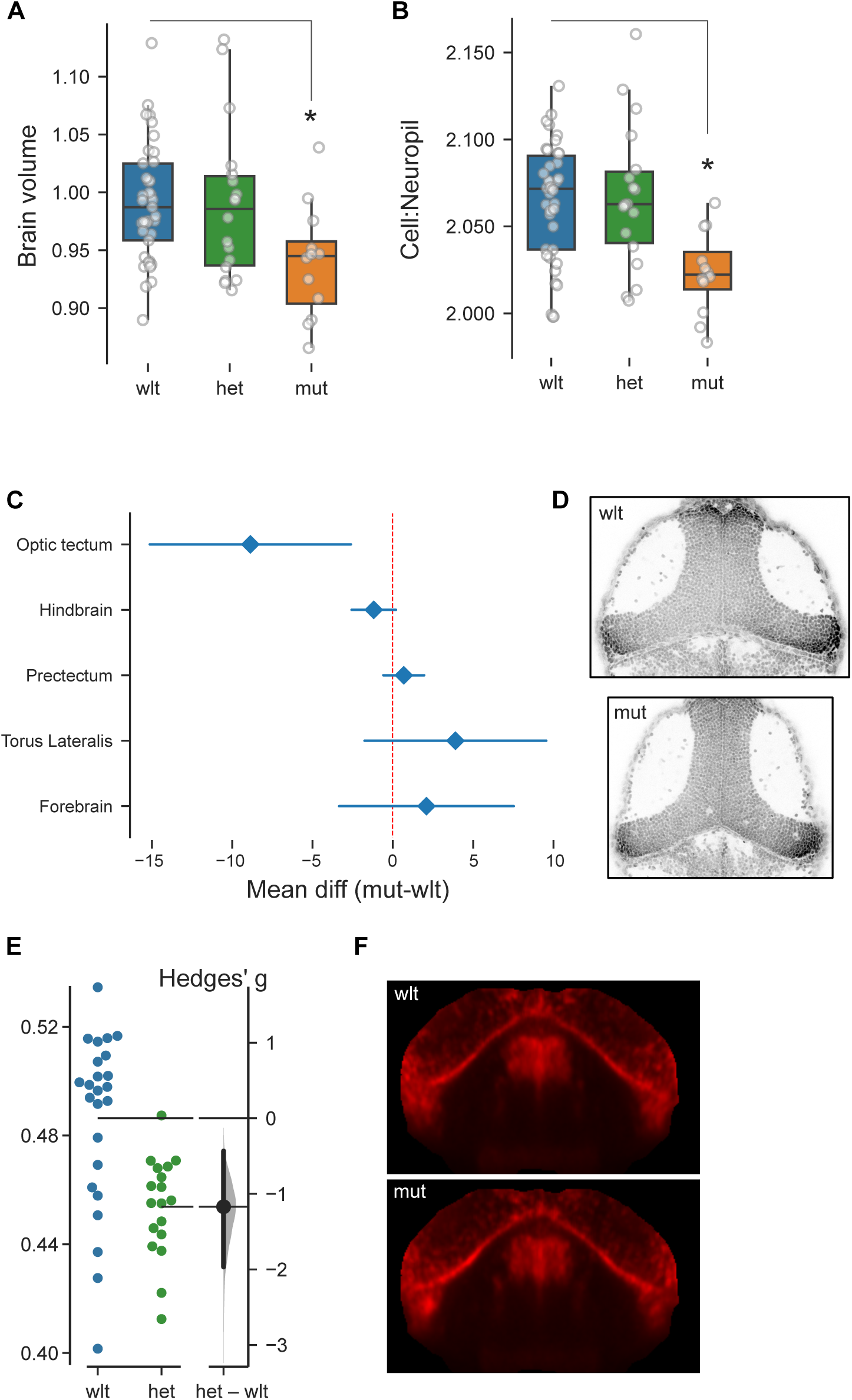
Whole-brain volume and cell density. A. Normalized brain volume in wild-type (N=37), heterozygous (N=18) and mutant (N=12) larvae. Each brain volume was normalized to the mean of the wild-type group. * t-test p < 0.01, comparing wild-type and mutants. B. Ratio of the volume of primarily cellular brain areas to neuropil dense areas same larvae as in (A). * t-test p < 0.001, comparing wild-type and mutants. C. Mean and confidence interval for difference in cell density for five brain regions in wild-type and mutant larvae (N=16,14 respectively), after DASPEI staining and manual cell counting. D. Hoechst33342 stained confocal section through the optic tectum in a wild-type and mutant larva. E. Volume of the midline cerebellar region that was reduced in TS2 homozygous larvae, in an independent cohort of wild-type (N=24) and sibling heterozygous (N=18) larvae. Mask volume normalized to total brain volume. F. Mean signal for *sox2* fluorescent *in situ* hybridization at the same coronal section as in Fig. 6D for wild-type (N=10) and mutant (N=9) TS2 larvae. Horizontal confocal scans were co-registered to ZBB, normalized, averaged and re-sliced for coronal view.

**Supplementary figure 7.**
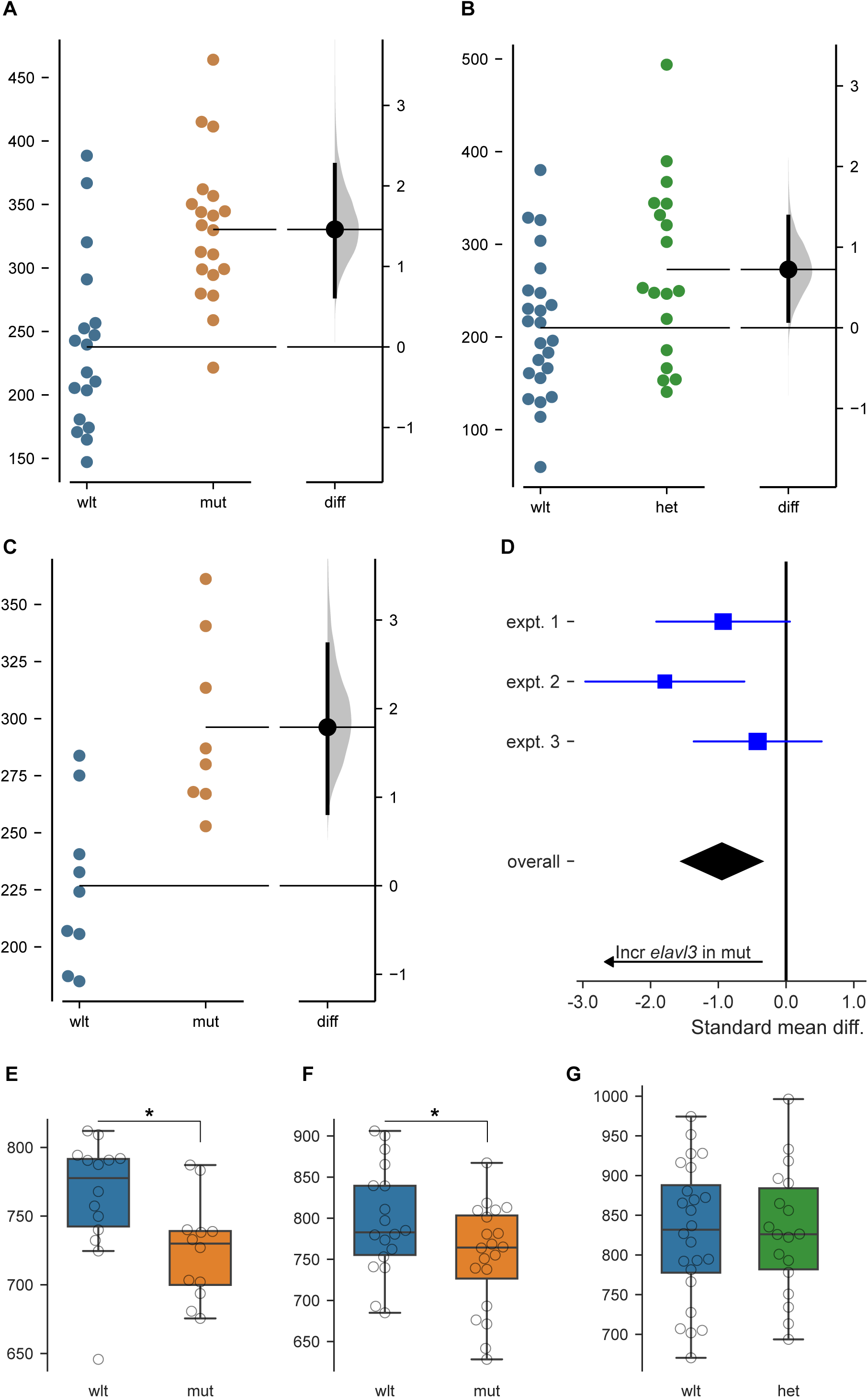
Distribution of post-mitotic neurons. A-B. Mean *tuba:mCardinal* signal within a mask encompassing part of the anterior hindbrain midline area with increased expression in Fig. 7A, in an independent cohort of TS2 wild-type and mutant larvae (N=18, 20 respectively; t-test, p < 0.001; A), and in wild-type and heterozygous larvae (N=24,18 respectively; t-test, p = 0.023; B). C. Midline *elavl3* fluorescent *in situ* hybridization signal intensity in the anterior hindbrain midline mask, in wild-type and mutant TS2 larvae (N=9 per genotype; t-test, p = 0.001). D. Meta-analysis of three experiments measuring anterior hindbrain midline *elavl3* signal in TS2 mutants and wild-type larvae. X-axis shows standard mean difference and confidence interval signal intensity between wild-type and mutant larvae in each experiment, and diamond shows a significant overall effect (standard mean difference −0.95, confidence interval [-1.54,-0.36]). There was moderate heterogeneity across experiments (I^2^ = 37.2%). E-G. Subpallial *gad1b:RFP* fluorescence intensity in TS2 wild-type and mutant larvae (E-F, independent cohorts) and wild-type versus heterozygous larvae (G). * t-test, p < 0.05.

**Supplementary figure 8.**
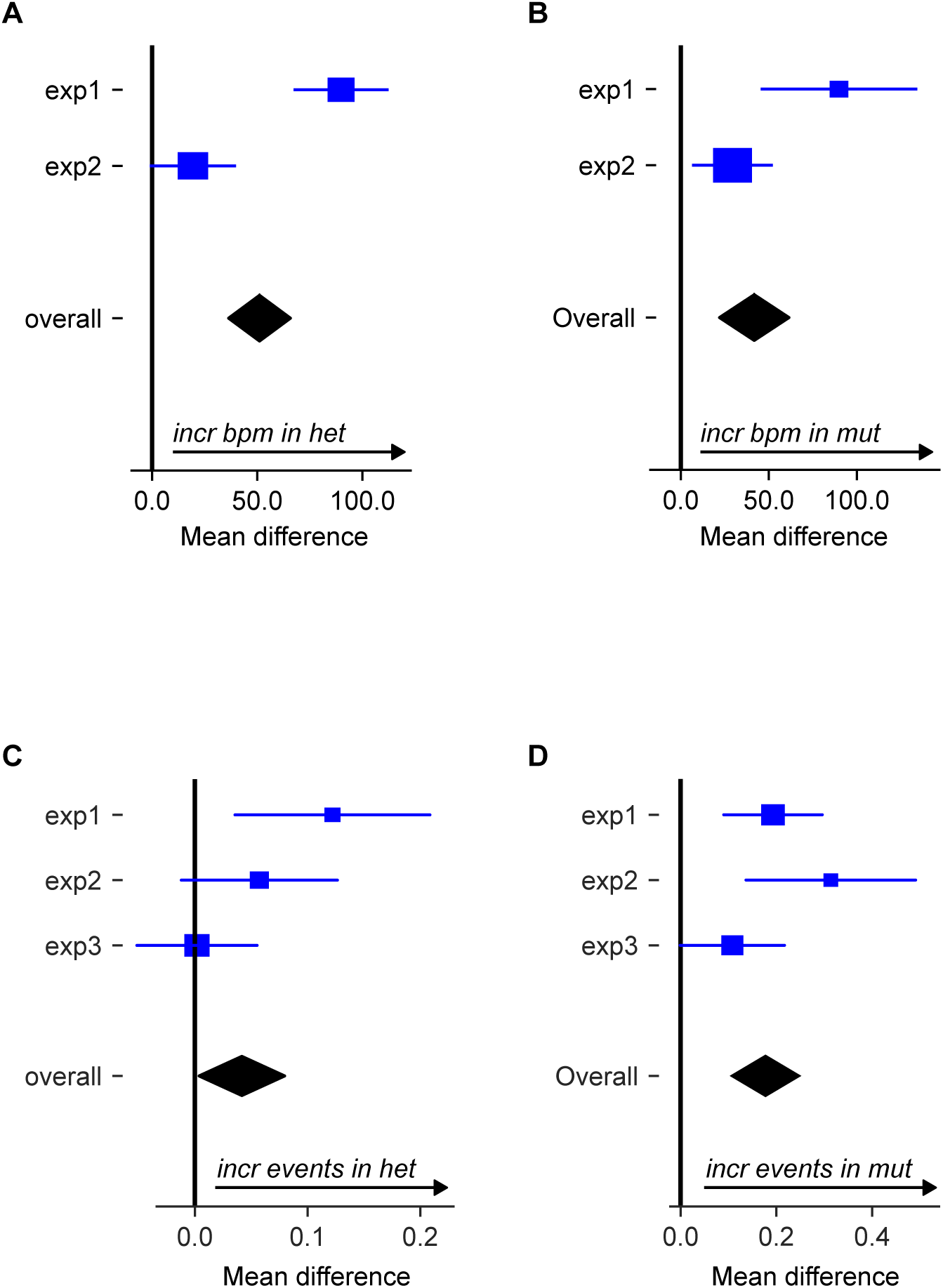
Tachycardia and seizure-like activity in heterozygous TS2 larvae at elevated temperature. A-B. Meta-analysis of two experiments measuring cardiac contraction frequency in TS2 wild-type and heterozygous (A) or mutant (B) sibling larvae. X-axis shows mean difference and confidence interval for difference between groups in each experiment. Diamond shows significant combined effect. A: mean difference 51 bpm, confidence interval [36.4,65.8], p < 0.001; B: mean difference 41.7 bpm, confidence interval [22.0,61.4], p < 0.001. C-D. Meta-analysis of three experiments measuring seizure-like behavior in *y680* wild-type and heterozygous (C) or mutant (D) sibling larvae. C: mean difference 0.04 events per minute, confidence interval [0,0.08], p=0.032. D: mean difference 0.18, confidence interval [0.11, 0.25], p < 0.001.

**Supplementary Table 1.**
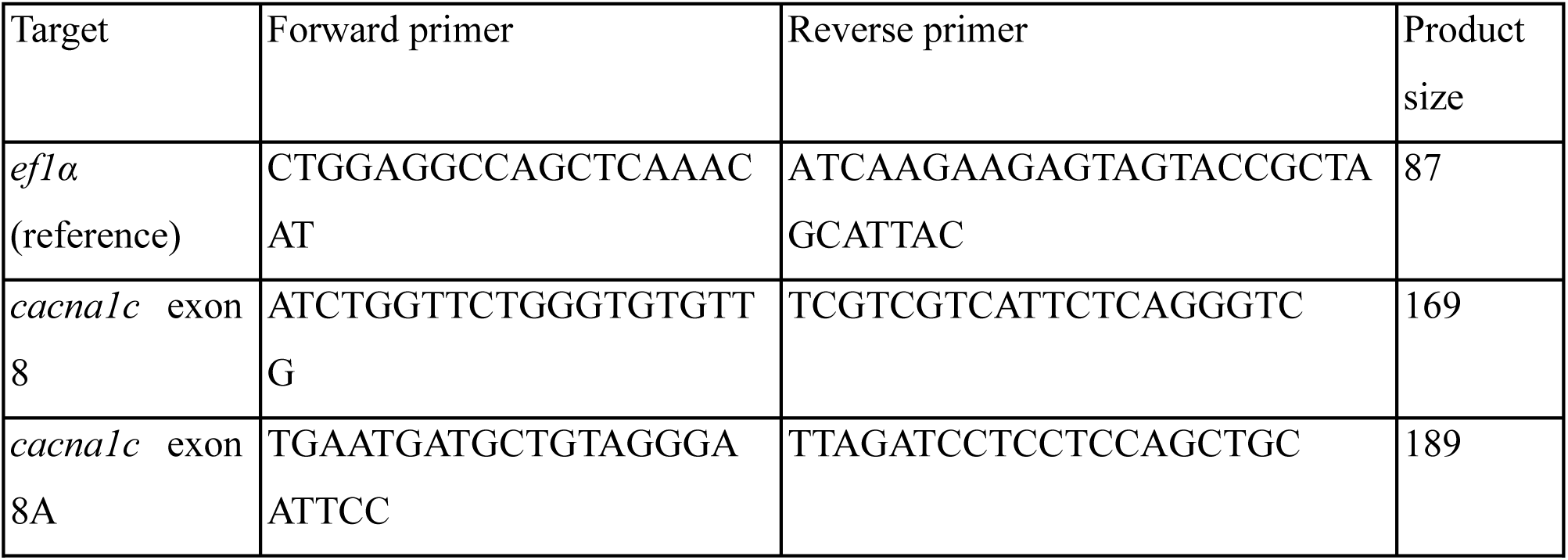
qPCR primers for measuring exon 8 and 8A usage in cacna1c.

## Notes

### Competing Interest Statement

The authors have declared no competing interest.

## References

1. R. Bauer, K. W. Timothy, A. Golden, Update on the Molecular Genetics of Timothy Syndrome. Front Pediatr 9, 668546 (2021).

2. G. Panagiotakos, C. Haveles, A. Arjun, R. Petrova, A. Rana, T. Portmann, S. P. Paşca, T. D. Palmer, R. E. Dolmetsch, Aberrant calcium channel splicing drives defects in cortical differentiation in Timothy syndrome. Elife 8, e51037 (2019).

3. Z. Z. Tang, S. Sharma, S. Zheng, G. Chawla, J. Nikolic, D. L. Black, Regulation of the mutually exclusive exons 8a and 8 in the CaV1.2 calcium channel transcript by polypyrimidine tract-binding protein. J Biol Chem 286, 10007–10016 (2011).

4. P. L. Bader, M. Faizi, L. H. Kim, S. F. Owen, M. R. Tadross, R. W. Alfa, G. C. L. Bett, R. W. Tsien, R. L. Rasmusson, M. Shamloo, Mouse model of Timothy syndrome recapitulates triad of autistic traits. Proc Natl Acad Sci U S A 108, 15432–15437 (2011).

5. S.-I. Horigane, Y. Ozawa, J. Zhang, H. Todoroki, P. Miao, A. Haijima, Y. Yanagawa, S. Ueda, S. Nakamura, M. Kakeyama, S. Takemoto-Kimura, A mouse model of Timothy syndrome exhibits altered social competitive dominance and inhibitory neuron development. FEBS Open Bio 10, 1436–1446 (2020).

6. F. Birey, J. Andersen, C. D. Makinson, S. Islam, W. Wei, N. Huber, H. C. Fan, K. R. C. Metzler, G. Panagiotakos, N. Thom, N. A. O’Rourke, L. M. Steinmetz, J. A. Bernstein, J. Hallmayer, J. R. Huguenard, S. P. Paşca, Assembly of functionally integrated human forebrain spheroids. Nature 545, 54–59 (2017).

7. F. Birey, M.-Y. Li, A. Gordon, M. V. Thete, A. M. Valencia, O. Revah, A. M. Paşca, D. H. Geschwind, S. P. Paşca, Dissecting the molecular basis of human interneuron migration in forebrain assembloids from Timothy syndrome. Cell Stem Cell 29, 248–264.e7 (2022).

8. X. Chen, F. Birey, M.-Y. Li, O. Revah, R. Levy, M. V. Thete, N. Reis, K. Kaganovsky, M. Onesto, N. Sakai, Z. Hudacova, J. Hao, X. Meng, S. Nishino, J. Huguenard, S. P. Pașca, Antisense oligonucleotide therapeutic approach for Timothy syndrome. Nature 628, 818–825 (2024).

9. M. Matsui, L. E. Lynch, I. Distefano, A. Galante, A. R. Gade, H.-G. Wang, N. Gómez-Banoy, P. Towers, D. S. Sinden, E. Q. Wei, A. S. Barnett, K. Johnson, R. Lima, A. Rubio-Navarro, A. K. Li, S. O. Marx, T. E. McGraw, P. S. Thornton, K. W. Timothy, J. C. Lo, G. S. Pitt, Multiple beta cell-independent mechanisms drive hypoglycemia in Timothy syndrome. Nat Commun 15, 8980 (2024).

10. J. C. Hocking, M. Distel, R. W. Köster, Studying cellular and subcellular dynamics in the developing zebrafish nervous system. Exp Neurol 242, 1–10 (2013).

11. T. Gupta, G. D. Marquart, E. J. Horstick, K. M. Tabor, S. Pajevic, H. A. Burgess, Morphometric analysis and neuroanatomical mapping of the zebrafish brain. Methods 150, 49–62 (2018).

12. W. Rottbauer, K. Baker, Z. G. Wo, M. A. Mohideen, H. F. Cantiello, M. C. Fishman, Growth and function of the embryonic heart depend upon the cardiac-specific L-type calcium channel alpha1 subunit. Dev Cell 1, 265–275 (2001).

13. K. V. Ramachandran, J. A. Hennessey, A. S. Barnett, X. Yin, H. A. Stadt, E. Foster, R. A. Shah, M. Yazawa, R. E. Dolmetsch, M. L. Kirby, G. S. Pitt, Calcium influx through L-type CaV1.2 Ca2+ channels regulates mandibular development. J Clin Invest 123, 1638–1646 (2013).

14. K. W. Timothy, R. Bauer, K. A. Larkin, E. P. Walsh, D. J. Abrams, C. Gonzalez Corcia, A. Valsamakis, G. S. Pitt, I. E. Dick, A. Golden, A Natural History Study of Timothy Syndrome. Orphanet J Rare Dis 19, 433 (2024).

15. D. Radzicki, H.-J. Yau, S. L. Pollema-Mays, L. Mlsna, K. Cho, S. Koh, M. Martina, Temperature-sensitive Cav1.2 calcium channels support intrinsic firing of pyramidal neurons and provide a target for the treatment of febrile seizures. J Neurosci 33, 9920–9931 (2013).

16. T. Nakajima, Y. Kaneko, M. Kurabayashi, Unveiling specific triggers and precipitating factors for fatal cardiac events in inherited arrhythmia syndromes. Circ J 79, 1185–1192 (2015).

17. A. Sur, Y. Wang, P. Capar, G. Margolin, M. K. Prochaska, J. A. Farrell, Single-cell analysis of shared signatures and transcriptional diversity during zebrafish development. Dev Cell 58, 3028–3047.e12 (2023).

18. M. Takeuchi, S. Yamaguchi, Y. Sakakibara, T. Hayashi, K. Matsuda, Y. Hara, C. Tanegashima, T. Shimizu, S. Kuraku, M. Hibi, Gene expression profiling of granule cells and Purkinje cells in the zebrafish cerebellum. J Comp Neurol 525, 1558–1585 (2017).

19. I. Splawski, K. W. Timothy, N. Decher, P. Kumar, F. B. Sachse, A. H. Beggs, M. C. Sanguinetti, M. T. Keating, Severe arrhythmia disorder caused by cardiac L-type calcium channel mutations. Proc Natl Acad Sci U S A 102, 8089–8096; discussion 8086-8088 (2005).

20. L. Yelin-Bekerman, I. Elbaz, A. Diber, D. Dahary, L. Gibbs-Bar, S. Alon, T. Lerer-Goldshtein, L. Appelbaum, Hypocretin neuron-specific transcriptome profiling identifies the sleep modulator Kcnh4a. Elife 4, e08638 (2015).

21. S. Thakker-Varia, J. J. Krol, J. Nettleton, P. M. Bilimoria, D. A. Bangasser, T. J. Shors, I. B. Black, J. Alder, The neuropeptide VGF produces antidepressant-like behavioral effects and enhances proliferation in the hippocampus. J Neurosci 27, 12156–12167 (2007).

22. S. Thakker-Varia, J. Behnke, D. Doobin, V. Dalal, K. Thakkar, F. Khadim, E. Wilson, A. Palmieri, H. Antila, T. Rantamaki, J. Alder, VGF (TLQP-62)-induced neurogenesis targets early phase neural progenitor cells in the adult hippocampus and requires glutamate and BDNF signaling. Stem Cell Res 12, 762–777 (2014).

23. T. Afrikanova, A.-S. K. Serruys, O. E. M. Buenafe, R. Clinckers, I. Smolders, P. A. M. de Witte, A. D. Crawford, C. V. Esguerra, Validation of the zebrafish pentylenetetrazol seizure model: locomotor versus electrographic responses to antiepileptic drugs. PLoS One 8, e54166 (2013).

24. G. D. Marquart, K. M. Tabor, M. Brown, J. L. Strykowski, G. K. Varshney, M. C. LaFave, T. Mueller, S. M. Burgess, S.-I. Higashijima, H. A. Burgess, A 3D Searchable Database of Transgenic Zebrafish Gal4 and Cre Lines for Functional Neuroanatomy Studies. Front Neural Circuits 9, 78 (2015).

25. G. D. Marquart, K. M. Tabor, E. J. Horstick, M. Brown, A. K. Geoca, N. F. Polys, D. D. Nogare, H. A. Burgess, High precision registration between zebrafish brain atlases using symmetric diffeomorphic normalization. Gigascience 6, 1–15 (2017).

26. V. Diep, L. H. Seaver, Long QT syndrome with craniofacial, digital, and neurologic features: Is it useful to distinguish between Timothy syndrome types 1 and 2? Am J Med Genet A 167A, 2780–2785 (2015).

27. A.-M. Haapanen-Saaristo, N. Virtanen, E. Tcarenkova, K. Vaparanta, M. Ampuja, E.-R. Vehniäinen, I. Paatero, Heat stress sensitizes zebrafish embryos to neurological and cardiac toxicity. Biochemical and Biophysical Research Communications 733, 150682 (2024).

28. B. W. Draper, P. A. Morcos, C. B. Kimmel, Inhibition of zebrafish fgf8 pre-mRNA splicing with morpholino oligos: a quantifiable method for gene knockdown. Genesis 30, 154–156 (2001).

29. P. L. Greer, M. E. Greenberg, From synapse to nucleus: calcium-dependent gene transcription in the control of synapse development and function. Neuron 59, 846–860 (2008).

30. A. Arjun McKinney, R. Petrova, G. Panagiotakos, Calcium and activity-dependent signaling in the developing cerebral cortex. Development 149, dev198853 (2022).

31. S. P. Paşca, T. Portmann, I. Voineagu, M. Yazawa, A. Shcheglovitov, A. M. Paşca, B. Cord, T. D. Palmer, S. Chikahisa, S. Nishino, J. A. Bernstein, J. Hallmayer, D. H. Geschwind, R. E. Dolmetsch, Using iPSC-derived neurons to uncover cellular phenotypes associated with Timothy syndrome. Nat Med 17, 1657–1662 (2011).

32. F. Zheng, X. Zhou, Y. Luo, H. Xiao, G. Wayman, H. Wang, Regulation of brain-derived neurotrophic factor exon IV transcription through calcium responsive elements in cortical neurons. PLoS One 6, e28441 (2011).

33. N. C. Spitzer, Electrical activity in early neuronal development. Nature 444, 707–712 (2006).

34. D. Bortone, F. Polleux, KCC2 expression promotes the termination of cortical interneuron migration in a voltage-sensitive calcium-dependent manner. Neuron 62, 53–71 (2009).

35. S. Kamijo, Y. Ishii, S.-I. Horigane, K. Suzuki, M. Ohkura, J. Nakai, H. Fujii, S. Takemoto-Kimura, H. Bito, A Critical Neurodevelopmental Role for L-Type Voltage-Gated Calcium Channels in Neurite Extension and Radial Migration. J Neurosci 38, 5551–5566 (2018).

36. D. P. Darcy, J. S. Isaacson, L-type calcium channels govern calcium signaling in migrating newborn neurons in the postnatal olfactory bulb. J Neurosci 29, 2510–2518 (2009).

37. W. Tang, J. D. Davidson, G. Zhang, K. E. Conen, J. Fang, F. Serluca, J. Li, X. Xiong, M. Coble, T. Tsai, G. Molind, C. H. Fawcett, E. Sanchez, P. Zhu, I. D. Couzin, M. C. Fishman, Genetic Control of Collective Behavior in Zebrafish. iScience 23, 100942 (2020).

38. J. Haverinen, M. Hassinen, S. N. Dash, M. Vornanen, Expression of calcium channel transcripts in the zebrafish heart: dominance of T-type channels. J Exp Biol 221, jeb179226 (2018).

39. E. Bovo, A. V. Dvornikov, S. R. Mazurek, P. P. de Tombe, A. V. Zima, Mechanisms of Ca^2^+ handling in zebrafish ventricular myocytes. Pflugers Arch 465, 1775–1784 (2013).

40. M. Vornanen, A. Badr, J. Haverinen, Cardiac arrhythmias in fish induced by natural and anthropogenic changes in environmental conditions. J Exp Biol 227, jeb247446 (2024).

41. C. Satou, Y. Kimura, S. Higashijima, Generation of multiple classes of V0 neurons in zebrafish spinal cord: progenitor heterogeneity and temporal control of neuronal diversity. J Neurosci 32, 1771– 83 (2012).

42. R. J. White, E. Mackay, S. W. Wilson, E. M. Busch-Nentwich, Allele-specific gene expression can underlie altered transcript abundance in zebrafish mutants. Elife 11, e72825 (2022).

43. S. Monti, P. Tamayo, J. Mesirov, T. Golub, Consensus Clustering: A Resampling-Based Method for Class Discovery and Visualization of Gene Expression Microarray Data. Machine Learning 52, 91–118 (2003).

44. S. X. Ge, D. Jung, R. Yao, ShinyGO: a graphical gene-set enrichment tool for animals and plants. Bioinformatics 36, 2628–2629 (2020).

45. Y. M. Bradford, C. E. Van Slyke, L. Ruzicka, A. Singer, A. Eagle, D. Fashena, D. G. Howe, K. Frazer, R. Martin, H. Paddock, C. Pich, S. Ramachandran, M. Westerfield, Zebrafish information network, the knowledgebase for Danio rerio research. Genetics 220, iyac016 (2022).

46. R. Tang, A. Dodd, D. Lai, W. C. McNabb, D. R. Love, Validation of zebrafish (Danio rerio) reference genes for quantitative real-time RT-PCR normalization. Acta Biochim Biophys Sin (Shanghai) 39, 384–90 (2007).

47. C. A. Schneider, W. S. Rasband, K. W. Eliceiri, NIH Image to ImageJ: 25 years of image analysis. Nature Methods 9, 671–675 (2012).

48. P. Virtanen, R. Gommers, T. E. Oliphant, M. Haberland, T. Reddy, D. Cournapeau, E. Burovski, P. Peterson, W. Weckesser, J. Bright, S. J. van der Walt, M. Brett, J. Wilson, K. J. Millman, N. Mayorov, A. R. J. Nelson, E. Jones, R. Kern, E. Larson, C. J. Carey, İ. Polat, Y. Feng, E. W. Moore, J. VanderPlas, D. Laxalde, J. Perktold, R. Cimrman, I. Henriksen, E. A. Quintero, C. R. Harris, A. M. Archibald, A. H. Ribeiro, F. Pedregosa, P. van Mulbregt, SciPy 1.0: fundamental algorithms for scientific computing in Python. Nat Methods 17, 261–272 (2020).

49. M. C. Halloran, M. Sato-Maeda, J. T. Warren, F. Su, Z. Lele, P. H. Krone, J. Y. Kuwada, W. Shoji, Laser-induced gene expression in specific cells of transgenic zebrafish. Development 127, 1953–60 (2000).

50. B. B. Avants, C. L. Epstein, M. Grossman, J. C. Gee, Symmetric diffeomorphic image registration with cross-correlation: evaluating automated labeling of elderly and neurodegenerative brain. Med Image Anal 12, 26–41 (2008).

51. A. A. Bhandiwad, T. Gupta, A. Subedi, V. Heigh, G. A. Holmes, H. A. Burgess, Brain Imaging and Registration in Larval Zebrafish. Methods Mol Biol 2707, 141–153 (2024).

52. J. Freeman, N. Vladimirov, T. Kawashima, Y. Mu, N. J. Sofroniew, D. V. Bennett, J. Rosen, C.-T. Yang, L. L. Looger, M. B. Ahrens, Mapping brain activity at scale with cluster computing. Nat Meth 11, 941–950 (2014).

53. E. Kuehn, D. S. Clausen, R. W. Null, B. M. Metzger, A. D. Willis, B. D. Özpolat, Segment number threshold determines juvenile onset of germline cluster expansion in Platynereis dumerilii. J Exp Zool B Mol Dev Evol 338, 225–240 (2022).

54. T. Yokogawa, M. C. Hannan, H. A. Burgess, The dorsal raphe modulates sensory responsiveness during arousal in zebrafish. J Neurosci 32, 15205–15215 (2012).

55. H. A. Burgess, M. Granato, Sensorimotor gating in larval zebrafish. J Neurosci 27, 4984–4994 (2007).

56. J. Ho, T. Tumkaya, S. Aryal, H. Choi, A. Claridge-Chang, Moving beyond P values: data analysis with estimation graphics. Nat Methods 16, 565–566 (2019).

57. M. Kunst, E. Laurell, N. Mokayes, A. Kramer, F. Kubo, A. M. Fernandes, D. Förster, M. Dal Maschio, H. Baier, A Cellular-Resolution Atlas of the Larval Zebrafish Brain. Neuron 103, 21–38.e5 (2019).

